# Binding of long non-coding RNAs within the androgen receptor N-terminal tail modulates interdomain communication and liquid-liquid phase separation

**DOI:** 10.64898/2026.05.28.728402

**Authors:** Nazanin Farahi, Galo Ezequiel Balatti, Alexander N. Volkov, Rita Pancsa, Peter Tompa, Remy Loris

## Abstract

The androgen receptor (AR) is a transcription factor whose overactivation is a primary driver of prostate cancer. Although AR interactions with several long non-coding RNAs (lncRNAs) have been implicated in castration-resistant prostate cancer, their underlying molecular mechanisms and functional consequences remain poorly understood. Here, we identify the N-terminal 37 residues of the intrinsically disordered AR N-terminal domain as the primary RNA-binding region that mediates selective interactions with the lncRNAs HOTAIR and SLNCR1. Residue-level mapping and mutational analysis define Y11, R13, and Q24 as key determinants of RNA recognition. This RNA-binding region partially overlaps with the F^23^QNLF^27^ motif, previously shown to mediate N/C interdomain communication with the ligand-binding domain through folding upon binding. We observe a partner-dependent binding mode in which this motif remains dynamically disordered upon RNA binding. LncRNA binding promotes phase separation of the N-terminal domain, indicating that lncRNAs upregulated in late-stage prostate cancer may lower the threshold for AR condensate formation and contribute to ligand-independent AR signaling. LncRNAs modulate communication between the N-terminal and ligand-binding domains within condensates, suggesting that lncRNA binding may tune hormone-dependent full-length AR signaling. These findings provide a mechanistic framework for AR-lncRNA regulation, laying the groundwork for future therapeutic strategies against advanced prostate cancer.

**Graphical abstract:** 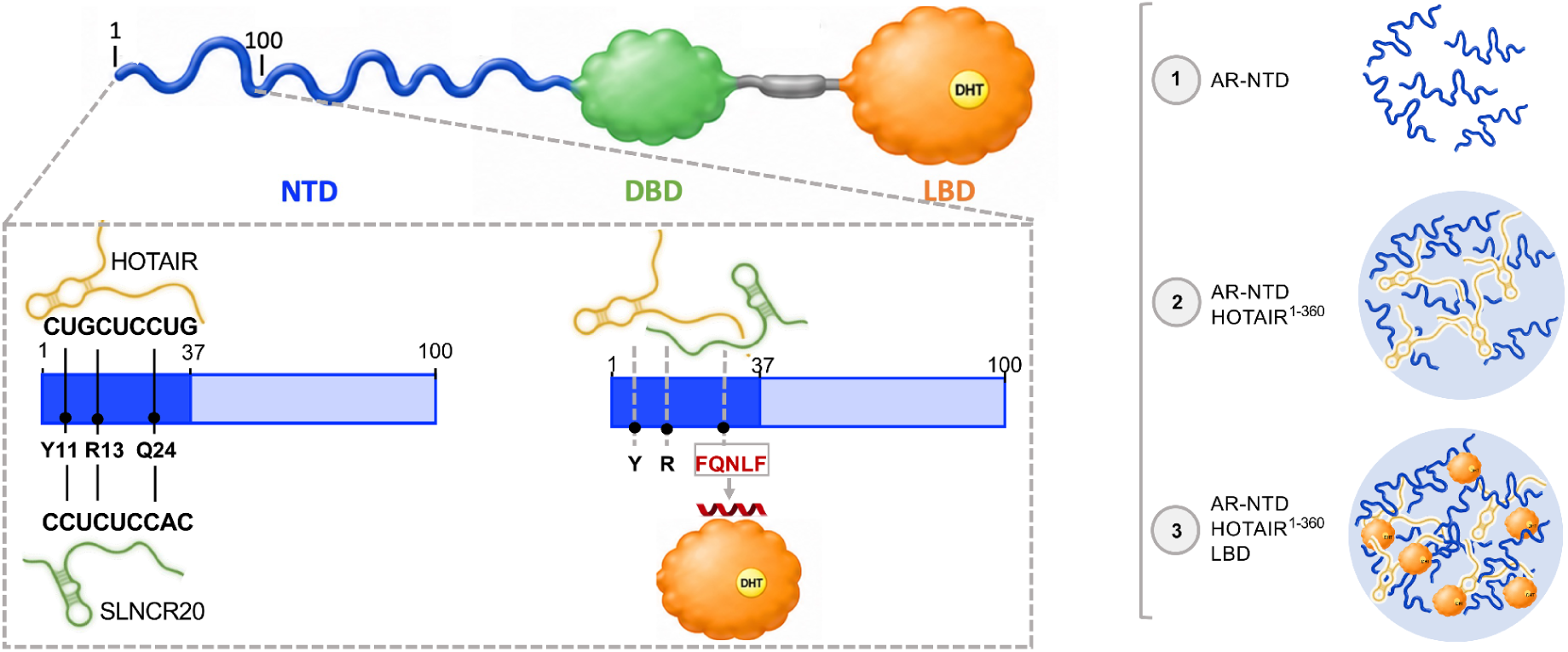

## Introduction

The androgen receptor (AR) plays a critical role in the development of prostate cancer, one of the most frequently diagnosed cancers in men globally (1). AR is a multidomain nuclear hormone receptor of 919 amino acids that comprises an intrinsically disordered N-terminal domain (NTD; hereafter AR^NTD^), followed by the folded DNA-binding domain and ligand-binding domain (LBD; hereafter AR^LBD^) (2). Its transcriptional activity is tightly regulated by androgens, making androgen deprivation therapy the cornerstone of prostate cancer treatment. Over time, however, prostate cancer overcomes androgen deprivation therapy, resulting in a relapse to the deadly form of the disease, androgen-independent castration-resistant prostate cancer (3,4). Drugs have been conventionally used to target the AR ligand-binding domain; however, these are often ineffective in late castration-resistant prostate cancer, primarily due to the emergence of splice variants (such as AR-V7) that lack the LBD and are therefore constitutively active (5).

Approximately a third of androgen receptor mutations identified in prostate cancer are located within the disordered N-terminal domain (residues 1-555; with no available X-ray structure (2)), indicating its direct involvement in disease pathogenesis (6). The NTD mediates interactions with diverse binding partners, such as the p160 family of AR coactivators, and also engages in intramolecular communication with the LBD (termed N/C interaction) via the F^23^QNLF^27^ motif, thereby modulating the transcriptional activity of the protein (7).

Importantly, among the diverse AR interactors, several long non-coding RNAs modulate AR function via poorly understood molecular mechanisms (8–11). Many AR-interacting lncRNAs, such as HOTAIR, PCAT1, SOCS2-AS1, LINC00675, and possibly PCGEM1 and PRNCR1, are upregulated in advanced prostate cancer cell lines, potentially contributing to androgen-independent AR activity and castration-resistant prostate cancer progression (12–17). Specifically, overexpression of HOTAIR was demonstrated to induce androgen-independent AR nuclear targeting and transcriptional activation (15).

Interestingly, AR activity upon androgen stimulation correlates with its ability to form or be recruited into transcriptional condensates at super-enhancers via liquid-liquid phase separation (LLPS), which is frequently deregulated in prostate cancer (18–22). The N-terminal domain of AR forms the primary determinant of AR phase separation, which is closely linked to its nuclear translocation and transactivation. Both full-length AR and the AR-V7 splice variant (which lacks the LBD) form nuclear condensates in cells, whereas the DNA-binding domain alone does not (18,20). AR^NTD^ phase separates in vitro at ∼100 μM in the presence of crowding agents (22) and together with SPOP E3 ubiquitin ligase through domain-motif interactions (21). Nevertheless, despite the dominant contribution of the NTD, all domains contribute to effective AR LLPS, as deleting the F^23^QNLF^27^ motif that mediates coupling between the NTD and LBD domains disrupts droplet formation in the full-length receptor (18,20).

AR-V7 undergoes phase separation less readily than AR-FL, requiring higher concentrations both in cells and in vitro, and its condensate-forming capacity is strongly dependent on cellular context, as it is observed in castration-resistant prostate cancer cell lines but only rarely in hormone-sensitive LNCaP cells (23). RNA molecules can promote LLPS by acting as multivalent co-scaffolds and lowering the effective protein concentration required for condensation (24,25); accordingly, many lncRNAs act as scaffolds for nuclear body assembly; for example, lncRNA DIGIT supports the formation of transcriptionally active ribonucleoprotein condensates (26). The ability of lncRNAs to bind AR N-terminal domain and modulate its chromatin occupancy suggests that they could also influence AR droplet dynamics at enhancer regions. Given that the expression of these lncRNAs is context-dependent and is normally upregulated in late-stage prostate cancer, we asked how they affect AR LLPS and whether their availability as interaction partners in castration-resistant prostate cancer cells may help explain the cell-type-specific LLPS behavior of AR-V7 (15).

Although many core aspects of AR function and prostate cancer pathogenesis have been thoroughly investigated, the highly interconnected regulatory mechanisms governing AR activity remain incompletely understood. lncRNAs have emerged as critical modulators within this regulatory landscape; however, the scope of their interactions with AR and their functional relevance remains poorly defined. Here, we study the molecular basis of AR-lncRNA regulation by mapping lncRNA-binding regions within the intrinsically disordered AR^NTD^ and answering fundamental questions regarding the interaction specificity, recognition principles, and underlying structural determinants. We further explore how lncRNA engagement influences NTD-driven LLPS of the androgen receptor, which is a critical process for understanding AR transcriptional regulation at super enhancers. Finally, we evaluate how these lncRNA interactions affect hormone-dependent N/C interdomain coupling in a reconstituted AR system.

## Materials and methods

### Proteins and nucleic acids

The amino acid sequence of human androgen receptor (Uniprot: P10275) was used as a template to generate all AR constructs used in this study. RNA oligonucleotides: SLNCR1-20nt (604-623 nucleotides of the RefSeq transcript NR_036488.1), HOTAIR-20nt (nucleotides 161-180 of the HOTAIR transcript annotation used in a previous study (15), equivalent to nucleotides 243-262 of the RefSeq transcript NR_186241.1), and the negative control RNA (RNA-NC-20nt) were ordered from Synpeptide. Unlabeled PolyU-20nt and PolyA-20nt oligos were purchased from Eurofins Genomics, while polyuridylic acid potassium salt (Poly U) was purchased from Sigma Aldrich. Yeast tRNA has been ordered from Thermo Fisher Scientific. An overview of the sequences of the different protein fragments and RNA oligonucleotides used in this study is provided in **Supplementary Table S1.**

The sequence of full-length HOTAIR was subcloned from LZRS-HOTAIR (AddGene plasmid #26110 (27)) into a pcDNA3.1 carrier vector downstream of the T7 promoter through HIFI assembly technique. The linear T7 promoter-containing DNA template of HOTAIR^1-360^ was then produced by PCR amplification of the region using linear_Hotair360_For and linear_Hotair360_Rev Primers **(Supplementary Table S2)**.

This PCR product was then transcribed in vitro using the HiScribe T7 High Yield RNA Synthesis Kit (New England Biolabs, NEB), according to the manufacturer’s instructions. Following transcription, the DNA template was removed by DNase I treatment, and the RNA product was purified using Monarch^®^ Spin RNA Cleanup Kit, quantified by UV absorbance at 260 nm, and stored at -80 °C until use.

### Cloning of AR protein constructs

The *Escherichia coli* codon-optimized coding sequence of human AR, containing a stop codon introduced after phenylalanine 554, was synthesized in a pET-21d(+) carrier plasmid downstream of a TEV cleavage site using EcoRI/XhoI restriction-ligation by GenScript. The T7NT*_MaSp_-AR expression plasmid **(Supplementary Figure 1)** was generated by cloning EcoRI/XhoI restriction-ligation of AR sequence into the multiple cloning site of an existing plasmid (pT7NT*_MaSp_-Aβ42, a gift from Dr. Henrik Biverstal, Karolinska Institutet; which was derived from the parent plasmid, pT7NT*-Bri2 113–231 R221E, Addgene plasmid #138134) (28,29). The resulting T7NT*_MaSp_-AR plasmid contains an N-terminal 6x His-tag followed by the NT*_MaSp_ solubility tag and the AR coding sequence, truncated at F554 by an introduced TAA stop codon, within its multiple cloning site under the T7 promoter. Site-directed mutagenesis was performed on the T7NT*_MaSp_-AR expression plasmid to generate two additional AR constructs, AR^1-100^ and AR^LBD^, using the primer sets listed in **Supplementary Table S2.** To generate the AR^NTD^ triple alanine mutant, site-directed mutagenesis was performed in a stepwise manner. The three substitutions were introduced stepwise, in the order of Q24A, Y11A, and R13A, and the final construct was verified by sequencing. The NTD^1-100^-3A mutant was subsequently generated from this full-length triple mutant construct using the NTD1-100 forward and reverse primers. The primers used are listed in **Supplementary Table S2**.

### Expression and purification of protein constructs

AR^NTD^ (1–554), AR^1-100^ (1–100), and their corresponding 3A mutants were expressed in *E. coli* BL21 Star™ (DE3) cells (NEB, Ipswich, MA, USA). Cells were grown at 37 °C in Luria Broth (LB) supplemented with 50 µg/mL Kanamycin until OD600 reached 0.6-0.8, after which protein expression was induced with 0.2 mM isopropyl β-d-1-thiogalactopyranoside (IPTG) and continued overnight at 16 °C with shaking at 175 rpm. Cells were harvested by centrifugation at 4,500 rpm for 20 min and resuspended in lysis buffers containing 20 mM TRIS-HCl pH 8.0, 500mM NaCl, 10 mM imidazole supplemented with 0.5 mM MgCl_2_, 0.1% Triton X-100, 1 µg/mL RNase A, 0.5 mM tris (2-carboxyethyl) phosphine (TCEP), 0.1 mM phenyl-methyl-sulfonyl fluoride (PMSF), 0.5 mM Benzamidine hydrochloride, and 1 of Roche complete protease inhibitor cocktail tablets. Cells were lysed on ice by sonication for 10 min (5 s pulse on, 5 s pulse off, 60% amplitude), followed by centrifugation at 20,000× *g* (4 °C) for 1 h in the presence of 1 mg/mL DNase I.

Since AR^NTD^ and its 3A mutant were recovered in the inclusion-body fraction, contaminating nucleic acids were removed using an adapted published washing protocol (30). Accordingly, the lysate supernatant was discarded, and the pellet was resuspended in 30 mL of 1× PBS pH 7.5, supplemented with the same additives, homogenized on ice, sonicated again using 5 s on/off pulses at 60% amplitude, and centrifuged at 20,000 × g for 45 min at 4 °C. Including this washing step reduced nonspecific nucleic acid contamination, resulting in an A260/A280 ratio of 0.58 in the final purified protein sample. The washed pellet was then solubilized in binding buffer containing 20 mM Tris-HCl, 1 M NaCl, 10 mM imidazole, 1 mM TCEP, and 8 M urea, pH 8.0, supplemented with the same additives. After sonication and centrifugation at 20,000 × g for 45 min at 4 °C, the clarified supernatant was filtered through a 0.45 µm filter and loaded onto a 5 mL nickel-charged HisTrap HP IMAC column. The column was washed with binding buffer containing 2 M NaCl, and the protein was eluted with 500 mM imidazole. The eluted sample was then buffer exchanged into TEV cleavage buffer containing 3 M urea, 200 mM NaCl, 50 mM sodium phosphate, and 1 mM TCEP, pH 7.0. To remove the NT*_MaSp_ solubility tag, the protein was incubated overnight at RT with in-house purified TEV protease at a 1:100 ratio. The following day, reverse IMAC was performed, and the cleaved AR-NTD protein was collected in the flow-through, while the His-tagged NT*_MaSp_ tag remained bound to the column and was recovered in the elution fraction. Protein in the flowthrough was concentrated and stored at -80 °C. On the day of the experiment, the sample was subjected to an additional size-exclusion chromatography (SEC) step to reduce aggregation and improve sample stability. SEC was performed using a Superdex 200 16/600 column pre-equilibrated with assay buffer containing 20 mM sodium phosphate, 200 mM NaCl, and 1 mM TCEP, pH 7.2.

AR^1-100^ and AR^1-100^-3A were recovered in the soluble fraction after lysis under native conditions. However, due to their tendency to aggregate and instability during later purification steps, the clarified supernatant was mixed with 2 volumes of 8 M urea-containing binding buffer, pH 7.0, and filtered through a 0.45 µm filter before loading onto a 5 mL HisTrap HP IMAC column. IMAC purification, buffer exchange, TEV cleavage, and reverse IMAC were performed as described for AR^NTD^, using the same buffers, except that the buffer pH was adjusted to 7.0 for these constructs. Following reverse IMAC, the cleaved protein was dialyzed into cation-exchange (CIEX) binding buffer containing 25 mM MES, 10 mM NaCl, 5 M urea, and 1 mM TCEP, pH 5.5. The protein was then captured by CIEX chromatography and eluted using a 1 M NaCl gradient. Fractions containing AR^1-100^ were concentrated and dialyzed overnight at 4 °C against storage buffer containing 20 mM sodium phosphate, 150 mM NaCl, 100mM KCl, and 1 mM TCEP, pH 7.2, and stored at -80 °C.

AR^LBD^ (residues 665-920) was expressed and lysed using the same procedure described for the AR^NTD^ constructs, with minor modifications. Briefly, prior to induction, cultures were cooled to 16 °C, supplemented with dihydrotestosterone (DHT) to a final concentration of 10 µM, and immediately induced overnight with 0.2 mM IPTG. Cells were harvested as described above, and the resulting pellets were stored at -80 °C. On the purification day, cell pellets were resuspended in IMAC binding/wash buffer containing 25 mM HEPES, 500 mM NaCl, 10% glycerol, 1 mM TCEP, 10 µM DHT, 1% Tween-20, and 20 mM imidazole, pH 7.4, supplemented with the same additives described above. Cells were lysed and centrifuged, and the soluble fraction was filtered before loading onto an IMAC column. Bound protein was eluted with 500 mM imidazole and desalted into TEV cleavage buffer containing 25 mM HEPES, 250 mM NaCl, 10% glycerol, 1 mM TCEP, 10 µM DHT, and 0.3% Tween-20, pH 7.4, using a HiPrep 26/10 desalting column. The NT*_MaSp_ tag was then cleaved overnight at 4 °C using TEV protease. The following day, reverse IMAC was performed, and the NaCl concentration was reduced to 100 mM through two 2-h dialysis steps. The cleaved AR^LBD^ was then captured by CIEX chromatography and eluted using a 1 M salt gradient. Finally, fractions containing pure AR^LBD^ were pooled and dialyzed overnight against storage buffer (25 mM HEPES, 200 mM NaCl, 10% glycerol, 1 mM TCEP, 10 µM DHT, and 1 mM EDTA, pH 7.2), and subsequently stored at -80.

### RNA and protein labeling

Short RNA oligonucleotides (SLNCR1-20nt, HOTAIR-20nt, PolyU-20nt, PolyA-20nt, RNA-NC-20nt) carrying Cy5 label at the 3’ end were ordered from Eurofins Genomics. PolyU and purified HOTAIR^1-360^ were 3′-end labeled with Cy5 or Cy3 using T4 RNA ligase and Cy5- or Cy3-labeled cytidine-5′-phosphate-3′-(6-aminohexyl)phosphate (triethylammonium salt) purchased from Jena Bioscience. AR^1-100^ and full-length NTD were labeled on lysine residues using Dylight® 488 fluorescent dye (Thermo Scientific) according to the manufacturer’s protocol. AR^LBD^ was labeled using the Monolith protein labeling kit RED-NHS 2nd Generation (NanoTemper Technologies).

### Electrophoretic mobility shift assay

Cy5-labeled SLNCR1-20nt RNA fragment at a final 50 nM concentration was mixed with the indicated amount of either recombinant AR^NTD^ or AR^1-100^ in 20 mM sodium phosphate, 20 mM KCl, 1 mM MgCl_2_, 0.5 mM TCEP, pH 7.2, supplemented with 2 μg yeast tRNA as a nonspecific RNA competitor. Binding reactions were assembled in a final volume of 20 μL and incubated in nuclease-free PCR tubes for 10 minutes at 4 degrees. After incubation, 3.5 µl of Ficoll 20% was added as loading solution to each sample before loading onto native polyacrylamide gels. AR^NTD^/RNA complexes were resolved on 6% native polyacrylamide gels prepared in 1X Sodium Citrate (10 mM sodium citrate, 20 mM NaCl, pH 5.8), whereas AR^1-100^/RNA complexes were resolved on 6% native polyacrylamide gels prepared in 1X TB buffer (89 mM Tris Base, 89 mM boric acid, adjusted to pH 7.2). Gels were pre-run at 60 V for 60 min at 4°C using pre-chilled running buffer. Electrophoresis was then performed at 110 V for 60 min at 4°C. Cy5-labeled RNA was detected by direct ingel fluorescence using the Odyssey M Imaging System (LI-COR Biosciences).

### Microscale thermophoresis

The Monolith™ NT.115 instrument (NanoTemper) was used to study the interaction of Cy5-labeled RNA fragments, including SLNCR1-20nt (20 nM), HOTAIR-20nt (20 nM), HOTAIR^1-360^ (50 nM), PolyU-20nt (22 nM), PolyA-20nt (22 nM), and RNA-NC-20nt (50 nM), with the indicated concentrations of recombinant AR^NTD^ or AR^1-100^ and their corresponding 3A mutant constructs. Experiments were performed either in a low salt buffer (20 mM sodium phosphate, pH 7.2, 20 mM KCl, 1 mM MgCl_2_, 1mM TCEP, 0.05% Tween-20) or a physiological salt buffer (20 mM sodium phosphate, pH 7.2, 20 mM KCl, 130 mM NaCl, 1 mM MgCl_2_, 1mM TCEP, 0.05% Tween-20) using Monolith™ NT.115 MST premium coated capillaries.

In case of N/C interdomain communication binding studies, 33 nM Cy5-labeled AR^LBD^ was titrated with increasing concentration of either AR^1-100^ or AR^1-100^-3A in buffer containing 20 mM sodium phosphate, pH 7.2, 200 mM NaCl, 1mM MgCl_2_, 0.5 mM TCEP, 10 μM DHT, 0.05% Tween-20. Reactions were then incubated for 10 minutes at 4 degrees prior to MST analysis. To probe how prior engagement of the AR^1-100^ by one binding partner influences subsequent binding of the other, AR^1-100^ dilution series were preincubated for 5 min at 4 °C with fixed concentrations of either binding partners 250 μM SLNCR1-20nt or 50 μM AR^LBD^, followed by addition of the Cy5-labeled third component AR^LBD^ (33 nM), or SLNCR1-20nt (20 nM), respectively. The reactions were further incubated for 10 min at 4 °C prior to MST analysis. All experiments were carried out at 50% MST power and 80% LED power. Equilibrium dissociation constants (Kd) were determined from triplicate measurements by fitting the data using MO. Affinity Analysis software.

### NMR spectroscopy

The samples were prepared in 20 mM sodium phosphate, pH 7.2, 20 mM KCl, 1 mM MgCl_2_, 0.5 mM TCEP, 0.02% NaN_3_, and 10% D_2_O for the lock. All NMR spectra were acquired at 293 K on a Bruker Avance III HD 800 MHz spectrometer, equipped with a TCI cryoprobe. The NMR data were processed with TopSpin 3.7 (Bruker) or NMRPipe (31) and analyzed in CCPNMR (31,32). Assignments of backbone amide resonances were taken from literature (33) and verified by 3D HNCACB, HN(CO)CACB, HNCO, and HN(CA)CO spectra acquired on a 1 mM sample of U-[^13^C,^15^N] AR NTD^1-100^.

The NMR binding experiments were performed by an incremental addition of SLNCR1-20nt or HOTAIR-20nt RNAs (5.0 mM or 6.1 mM stock solutions, respectively) to 0.2 mM samples of U-[^13^C,^15^N] AR^1-100^, with the spectral changes monitored in [^1^H,^15^N] HSQC spectra acquired at each increment. The backbone amide resonance of H101, which showed substantial chemical shift variability across protein batches, was excluded from the analysis. The average chemical shift perturbations (Δδ_avg_) were calculated as Δδ_avg_ = (Δδ_N_ ^2^/50 + Δδ_H_ ^2^/2)^0.5^, where Δδ and Δδ are the chemical shift changes of the backbone amide nitrogens and protons, respectively. The NMR titration curves were analyzed with a two-parameter nonlinear least squares fit using a one-site binding model corrected for the dilution effect (34), Equation 1:

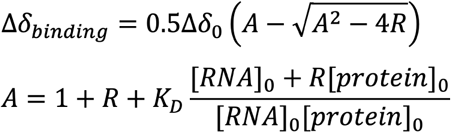

where Δδ_binding_ is the chemical shift perturbation at a given RNA/protein ratio; Δδ_0_ is the chemical shift perturbation at 100 % protein bound; R is the [RNA]/[protein] ratio at a given point; [protein]_0_ and [RNA]_0_ are the concentrations of the starting protein sample and the RNA titrant stock solution, respectively; and Kd is the equilibrium dissociation constant.

### Fluorescence microscopy of in vitro protein condensation

Droplet images were acquired using a Leica DMi8 microscope connected to a Leica DFC7000 GT camera. AR^NTD^ and AR^NTD^-3A protein samples were prepared by adding approximately 1% of Dylight® 488-labeled corresponding protein and kept on ice. Samples were assembled by mixing the protein solution at the indicated protein concentration with 20 mM sodium phosphate, pH 7.2, 200 mM NaCl, 1 mM MgCl_2_, and 0.5 mM TCEP buffer (for experiments including AR^LBD^, the buffer was also supplemented with 10 μM DHT required for its solubility). Unlabeled RNA partners, including PolyU, HOTAIR-20nt, SLNCR1-20nt, and HOTAIR^1-360^ or AR^LBD^ protein, were then added in excess at determined concentrations together with their corresponding fluorescently labeled molecules. Reactions were incubated for 2 minutes at RT in non-binding black 384-well plates with transparent bottom (Greinerbio-one,μClear®). Subsequently, 10 μl of each sample was transferred to a glass slide, covered with a coverslip, and imaged immediately using a ×100/1.4 Oil-Immersion objective lens. Since PolyU is heterogeneous in length, its concentration cannot be precisely defined based on polymer-chain molarity. Therefore, it was added at a nucleotide-residue amount equivalent to 25 μM HOTAIR^1-360^ rather than on a polymer-chain molarity basis. In a 15 μL reaction, this corresponded to 135 nmol uridylate residues, equivalent to ∼41.3 μg PolyU.

For the RNA and AR^LBD^ enrichment and colocalization analysis, AR^NTD^-containing droplets were first generated by incubating AR^NTD^ at the indicated concentration with either equimolar HOTAIR^1-360^ or AR^LBD^, together with a small fraction of the corresponding fluorescently labeled proteins. After 2 min of incubation, the second binding partner was added at the indicated final concentration, and samples were incubated for an additional 3 min before imaging. Images were acquired in the relevant fluorescence channels using identical microscope settings for all directly compared conditions, and raw images were used for the quantitative analyses.

### Fluorescence recovery after photobleaching (FRAP)

Sample preparation was identical to that described above for the microscopy experiments. For FRAP measurements, 10 pre-bleach images were acquired at 50 ms intervals, followed by 50 ms of photobleaching using the 488 nm laser at 100% power. Post-bleach images were then recorded every second for 2 min to monitor fluorescence recovery.

### Image analysis

#### In vitro droplet fluorescence quantification

For fluorescence quantification in **Figure 9**, droplets were defined as regions of interest (ROIs) in the AR^NTD^ channel, and corresponding ROIs were transferred to the RNA or AR^LBD^ channel. Mean fluorescence intensity was measured for each droplet in three regions: inside the droplet, in the surrounding dilute phase, and in a local background region. Enrichment of the fluorescent binding partner was calculated for each droplet as a background-corrected partition coefficient, Kp = (I^inside^−I^bg^)/(I^outside^−I^bg^), providing a measure of dense-phase versus dilute-phase enrichment. Image analysis was performed in Fiji/ImageJ. Dots in **Figure 9** represent individual droplets pooled from 10 images across three biological replicates. Droplet distributions were compared using an unpaired Mann-Whitney test.

### Line-scan colocalization analysis

Representative line-scan intensity profiles were generated in Fiji/ImageJ from raw multichannel fluorescence images. A fixed-width line ROI was drawn through the center of a representative droplet in the AR^NTD^ channel and applied to the corresponding partner channel. Fluorescence intensity profiles were extracted using the Plot Profile function, normalized to the maximum intensity in each channel, and plotted against distance (μm). The shown profile is representative of the same distribution pattern observed across multiple droplets in 3 biological replicates.

### Bioinformatics analysis

#### AR^NTD^ charge distribution and RNA binding site(s) predictions

DisoRDPbind (35) and Bindembed21 (36) RNA-binding site prediction tools were used via their webserver interface to predict residue-level RNA-binding propensity within AR^NTD^. Residues with DisoRDPbind propensity score ≥ 0.151 were considered as putative RNA-binding sites according to the default threshold recommended by the webserver. However, for Bindembed21 (36) predictions, a 0.2 cutoff was used only as a permissive, hypothesis-generating threshold to identify candidate interaction regions, which were subsequently tested experimentally.

A previously published charge-based cross-correlation method was used to identify similar HIV Tat ARM-like sequences within AR^NTD^ (37). Accordingly, the Tat ARM sequence “RKKRRQRRR” was digitized to the amino acid charge pattern “111110111” to create a 9-mer search kernel. The AR^NTD^ sequence was then digitized to “1” for R/K amino acid residues and “0” otherwise. NTD sequence was further refined by setting all entries to “0” in 9-mer windows where no R’s were present. The cross-correlation between the search kernel and the NTD sequence was then computed using the ‘correlate’ function in Scipy, with the “direct” method.

AR^NTD^ charge distribution at pH 7.4, with a 20-aa sliding-window smoothing, was plotted using VOLPES (38) (https://volpes.lisc.univie.ac.at/).

### Motif search in AR-interacting lncRNAs

To identify SLNCR1-like sequence motifs in AR-binding lncRNAs, full-length sequences of HOTAIR, SLNCR1, PRNCR1, SOCS2-AS1, PCAT1, PCGEM1, and LINC00675 were retrieved from the NCBI database. Motif scanning was performed using FIMO from the MEME Suite (v5.5.9) with the consensus 9-nt SLNCR1 motif, CYUCUCCWS, as the query. Uniform background nucleotide frequencies were applied (A = 0.25, C = 0.25, G = 0.25, U = 0.25). The position-specific probability matrix used for scanning was as follows:

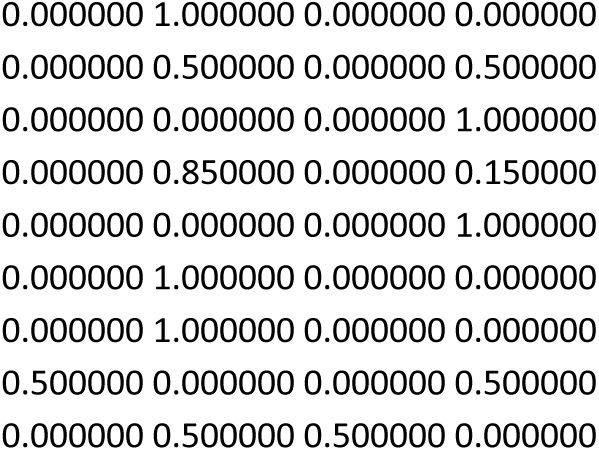

In transcripts for which AR-interacting regions had been experimentally defined (SLNCR1^568–637^, HOTAIR^1–360^, and PCGEM^421–480^, LINC00675^737-1530^) (9,12,15,17), the search was confined to those relevant regions. For lncRNAs lacking prior regional information, full-length transcripts were analyzed using FIMO, applying a significance threshold of *p* < 0.001 in all cases except PCGEM1, for which a more lenient threshold of *p* < 0.01 was used. Under these criteria, 36 motif occurrences were identified across the seven lncRNAs. Candidate motifs were then prioritized based on statistical support, selecting the site or sites with the lowest *p*-value. The final set of FIMO-identified motifs, together with the candidate motifs selected for experimental follow-up, is listed in **Supplementary Table S3.**

### Molecular dynamics simulations

Initial coordinates for the AR N-terminal fragment AR^1-100^ were obtained from AlphaFold (39) (entry AF-P10275-F1). The tertiary structures of the RNA fragments HOTAIR-20nt, SLNCR1-20nt, and PolyA-20nt negative control were predicted using RNAComposer (40).

All-atom MD simulations were performed for AR^1-100^ in the presence of HOTAIR-20nt, SLNCR1-20nt, or PolyA-20nt. Unbiased simulations were initiated from non-preformed complexes, with protein and RNA placed in spatially separated configurations in bulk solvent. No restraints, docking constraints, or predefined contacts were imposed to favor binding, allowing RNA association with AR^1-100^ to emerge spontaneously during the trajectories. Each unbiased production simulation was extended to 2 μs. In parallel, independent HADDOCK-derived MD refinement simulations were performed for the AR-HOTAIR-20nt and AR-SLNCR1-20nt systems. For each RNA fragment, the top-ranked HADDOCK model was used as the starting structure and simulated for 1 μs to assess the stability of docking-compatible interfaces and their consistency with interaction regions sampled in the unbiased simulations.

Simulations were carried out in GROMACS (59) using the CHARMM36m force field (60) for the protein and compatible parameters for RNA and ions. CHARMM36m was selected because it improves the description of both folded and intrinsically disordered proteins, which is relevant for AR^1-100^, a region combining extensive disorder with short linear motifs involved in partner recognition, including the RNA-binding region and the FQNLF-containing segment. CHARMM36/CHARMM36m force fields have also been benchmarked in peptide self-assembly systems, including Aβ16-22, where they provide a reasonable balance between secondary-structure formation and assembly kinetics compared with alternative atomistic force fields (61). Thus, a CHARMM36m/CHARMM-compatible setup was used to describe both the dynamic AR^NTD^ ensemble and its interactions with RNA fragments in an internally consistent all-atom framework.

Each system was solvated with the CHARMM-modified TIP3P water model, neutralized with Na⁺ and Cl⁻ ions, and adjusted to a final NaCl concentration of 0.15 M. Electrostatic interactions during production simulations were treated using the particle-mesh Ewald (PME) method, with a 1.2 nm cutoff for short-range interactions. Lennard-Jones interactions were smoothly switched off between 1.0 and 1.2 nm. All bonds involving hydrogen atoms were constrained using LINCS, allowing a 2 fs integration time step. After energy minimization, systems were equilibrated stepwise, starting from low temperature under positional restraints, followed by gradual heating to 303.15 K and progressive release of backbone, side-chain, and dihedral restraints. Pressure coupling was introduced during intermediate equilibration stages to stabilize system density. Production simulations were performed in the NPT ensemble at 303.15 K and 1 bar, using the Nosé-Hoover thermostat and Parrinello-Rahman barostat. Each production trajectory was extended up to 2 μs, unless otherwise indicated.

Trajectory analyses were performed using GROMACS tools: *gmx energy* for interaction-energy calculations, *gmx distance* for residue-RNA distances and time-resolved contacts, and *gmx mdmat* for mean minimum-distance contact maps. For interaction-energy analyses, Coulomb and Lennard-Jones contributions between AR^1-100^ and RNA fragments were calculated using a reaction-field setup to assess the relative contributions of electrostatic and short-range non-electrostatic interactions during recognition and stabilization. Mean minimum-distance contact maps were computed over defined trajectory windows to identify persistent RNA-contacting regions on AR^1-100^.

Residue-level protein-RNA interactions were characterized using the Residue Interaction Network Generator (RING (41)), which identifies noncovalent contacts from atomic coordinates based on geometric and chemical criteria. RING was applied to MD trajectory ensembles to classify recurrent interactions, including hydrogen bonds, van der Waals contacts, π-π stacking, cation-π, ionic, and π-hydrogen interactions, and to estimate their occupancies across the simulations. Strict geometric thresholds were used (hydrogen bond donor–acceptor ≤ 3.9 Å, H–acceptor ≤ 2.5 Å, π–π stacking ≤ 6.5 Å, cation–π ≤ 5 Å, ionic ≤ 4 Å, π–hydrogen ≤ 4.3 Å), and only interactions with occupancies above 20% were retained for analysis and reported in **Supplementary Tables S4 and S5**.

Independent HADDOCK (42) docking calculations were also performed using the same initial protein and RNA structures, focusing on the N-terminal 1-40 region of AR^1-100^. HADDOCK is an information-driven flexible docking approach that incorporates prior knowledge of interaction interfaces through ambiguous interaction restraints (AIRs), enabling the modeling of protein-nucleic acid complexes beyond purely ab initio strategies. For each system, the top-ranked HADDOCK model was selected for subsequent short MD refinements to assess the stability and consistency of the predicted RNA-protein interfaces.

## Results

### The RNA-binding region of AR is located in the first 37 amino acids of its N-terminal domain

Previous studies showed that the disordered N-terminal region of AR interacts directly with lncRNAs such as HOTAIR, SLNCR1, LINC00675, and PCGEM1 (12,15,17,43). Predictions using DisoRDPbind (35), a predictor of disordered RNA-binding regions, suggest that the first 37 residues constitute an RNA-binding region (**Figure 1A**). This region (hereafter referred to as AR^1-37^) exhibits a net positive charge, making it ideal for binding to the negatively charged RNAs (**Figure 1B**). The same region is further supported by BindEmbed21 (36) predictions showing a modest but spatially clustered increase in nucleic-acid-binding propensity compared to the rest of the N-terminal domain sequence (**Figure 1C**). Finally, charge-based cross-correlation with the HIV Tat arginine-rich motif (ARM) RKKRRQRRR identifies a moderate ARM-like charge pattern within AR^1-37^ corresponding to the ^9^RVYPRPPSKTYRG^21^ sequence (**Figure 1D**), in line with a recent study suggesting that at least half of the transcription factors use similar basic motifs within disordered regions to bind RNA (37). Together, these independent sequence-based analyses nominate AR^1-37^ as a positively charged, RNA-binding-prone region within the AR N-terminal domain.

**Figure 1.**
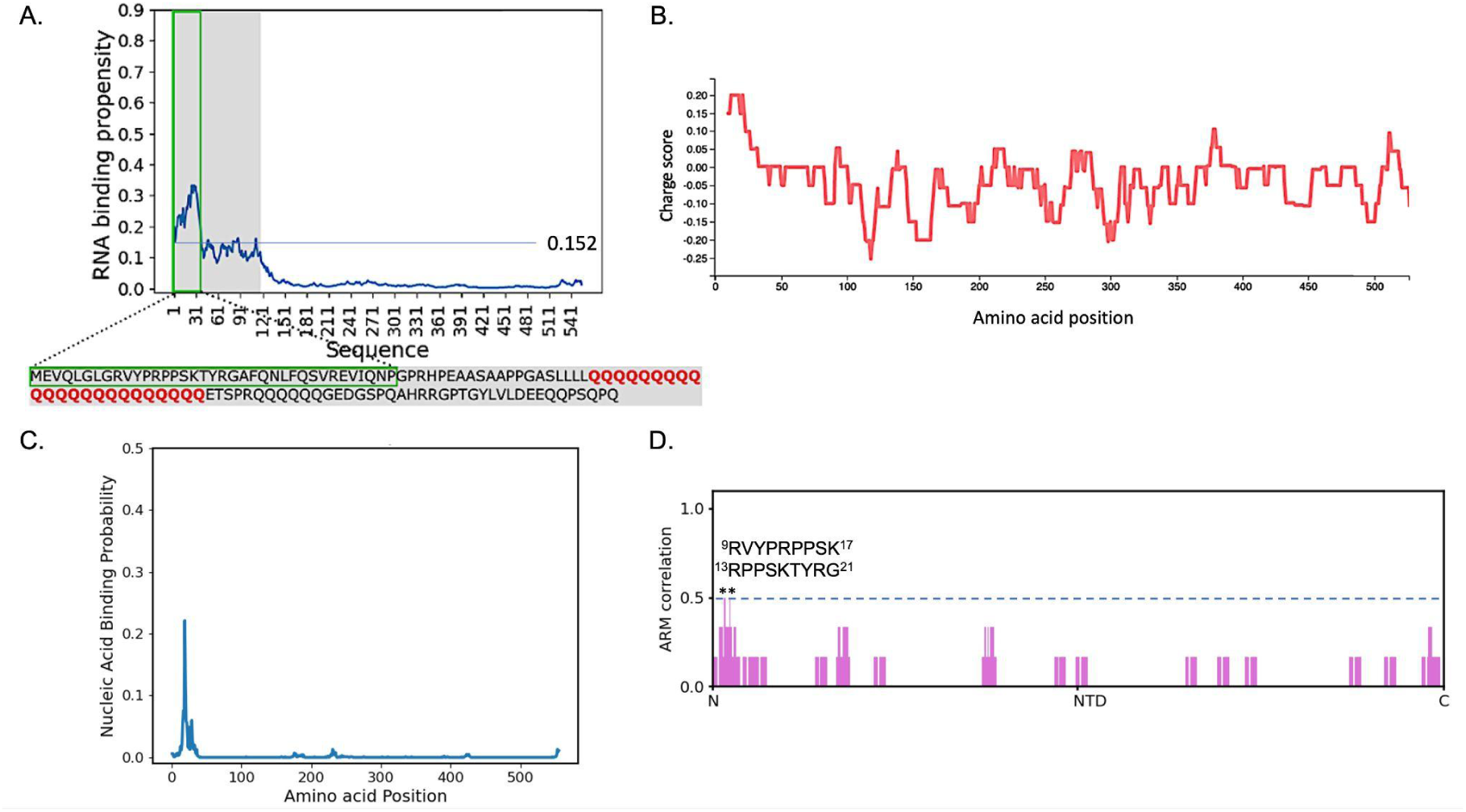
Sequence-based predictions point to a potential RNA-binding region within the AR N-terminal domain. A) DisoRDPbind RNA binding prediction for AR^NTD^. The residues with the highest RNA-binding propensity (residues 1-37) are highlighted in green. B) AR^NTD^ charge distribution at pH 7.4 with a 20-aa sliding window smoothing generated using VOLPES. C) BindEmbed21 predictor predicts the N-terminal region of NTD as a putative NA-binding site. D) Diagram of AR^NTD^ and its charge-based cross-correlation profile, highlighting regions enriched in R/K motifs across a sliding window (*highest-scoring ARM-like region).

To experimentally test whether the predicted N-terminal RNA-binding region mediates lncRNA recognition, we focused on SLNCR1 and HOTAIR lncRNAs. SLNCR1 was selected because prior work identified a pyrimidine-rich motif, “C[CU]U[CU]UCC[AU][GC]”, that mediates its binding to the AR^NTD^ (43). HOTAIR was selected as the second, prostate cancer-relevant candidate based on clinical evidence of patient samples and transcriptomic data showing its elevated expression in metastatic prostate cancer, association with poor clinical outcomes, and therapeutic resistance (15,44,45). To map the interaction site(s) within AR^1-37^, we used RNA fragments previously implicated in the binding of AR^NTD^: nucleotides 604-623 of SLNCR1 (hereafter referred to as SLNCR1-20nt) and nucleotides 1-360 of HOTAIR (HOTAIR^1-360^) (15,43). Electrophoretic mobility shift assays (EMSAs) reveal concentration-dependent complex formation between SLNCR1-20nt and both AR^NTD^ and its N-terminal segment encompassing the first 100 residues (AR^1-100^) (**Figure 2A, B**). Microscale thermophoresis (MST) measurements nevertheless show a weaker binding affinity of SLNCR1-20nt for AR^NTD^ compared to AR^1-100^, with dissociation constants of 284 ± 78.4 µM and 36 ± 4.3 µM, respectively (**Figure 2C**). MST experiments equally establish AR^1-100^ as the core binding region for HOTAIR^1-360^, with an affinity of approximately 50 µM (**Figure 2D**). Collectively, these results demonstrate that AR^1-100^ acts as the principal RNA-binding region, while distal regions within the intrinsically disordered AR^NTD^ apparently attenuate binding.

**Figure 2.**
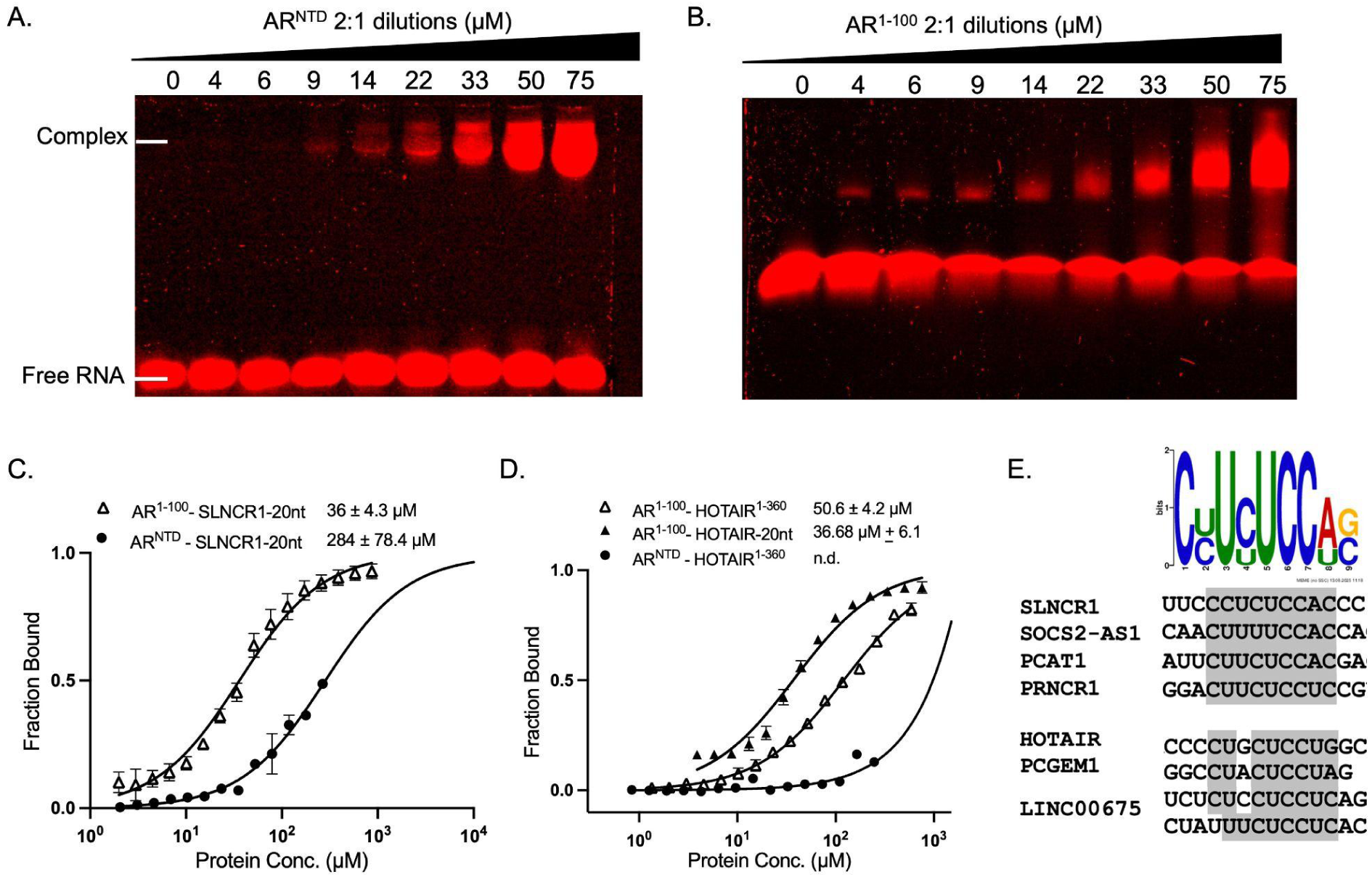
The N-terminal 100-residue segment of AR^NTD^ is sufficient to mediate interaction with lncRNAs. A, B) REMSA analysis of 50 nM Cy5-labeled SLNCR1-20nt incubated with increasing concentrations of AR^1-100^ and AR^NTD^. Bands correspond to the free labeled RNA probe and the more slowly migrating RNA-protein complexes, resolved on a 6% TB gel for AR^1-100^ and a 6% Sodium Citrate gel for full-length NTD. C) MST dose-response curves for the binding interaction between 20 nM Cy5-labeled SLNCR1-20nt and increasing concentrations of indicated NTD constructs, with Kd values indicated. Error bars represent the standard deviation (SD) of three biological replicates. D) MST dose-response curves for the binding of 50 nM Cy5-labeled HOTAIR^1-360^ or 20 nM Cy5-labeled HOTAIR-20nt with indicated NTD constructs, along with corresponding Kd values. Error bars represent the SD of three biological replicates. All binding assays were performed in 20 mM NaPi, pH 7.2, 20 mM KCl, 1 mM MgCl_2_, 1 mM TCEP, 0.05% Tween-20. E) Consensus and near-consensus motifs identified within AR-binding lncRNA sequences are indicated by gray boxes.

### HOTAIR^1-360^ interacts with AR via a near-consensus CUGCUCCUG motif

We next examined whether any conserved sequence motifs could explain the shared ability of AR-associated lncRNAs to bind the AR^NTD^ region. To address this, we focused on the recently defined SLNCR1 motif “C[CU]U[CU]UCC[AU][GC]” (43). Using FIMO motif analysis (46), we find consensus or near-consensus instances of this motif in two groups of AR-associated lncRNAs that are upregulated in castration-resistant prostate cancer cell lines, respectively: (1) SOCS2-AS1, PCAT1, and PRNCR1, and (2) HOTAIR, PCGEM1, and LINC00675 (**Figure 2E; see Materials and methods**).

Within HOTAIR^1-360^, the sequence CUGCUCCUG (nucleotides 167-175) closely matches the AR-binding consensus. To test whether this region mediates AR^NTD^ binding, we performed MST using a 20-nt HOTAIR fragment spanning nucleotides 161-180 (referred to as HOTAIR-20nt), which binds NTD^1-100^ with a Kd of ∼ 36 µM **(Figure 2D)**. This is essentially identical to the affinity of AR^1-100^ for SLNCR1-20nt and comparable to the affinity of the longer HOTAIR^1-360^ fragment (∼50.6 ± 4.2 µM; **Figure 2D**).

### Distinct molecular driving forces underlie specific AR interactions with HOTAIR-20nt and SLNCR1-20nt

To assess the specificity of AR^1-100^ for the C[CU]U[CU]UCC[AU][GC] RNA motif, the binding of SLNCR1-20nt and HOTAIR-20nt (**Figure 3A**) was compared to nonspecific 20-nt PolyA and PolyU RNAs (PolyA-20nt and PolyU-20nt). While SLNCR-20nt and HOTAIR-20nt showed clear binding to AR^1-100^ in EMSA experiments, no complex formation is observed for PolyA-20nt or PolyU-20nt (**Figure 3B**). These findings indicate that the interaction is selective and not solely driven by nonspecific electrostatic interactions with generic RNA. Consistent with our EMSA results, MST confirms that both Cy5-labeled SLNCR1-20nt and HOTAIR-20nt bind AR^1-100^ with similar affinities of approximately 36 µM. In contrast, AR^1-100^ does not bind Cy5-labeled PolyA-20nt or PolyU-20nt and shows substantially weaker binding to the length- and composition-matched negative control RNA, RNA-NC-20nt, with a Kd of ∼600 µM **(Figure 3C)**.

**Figure 3.**
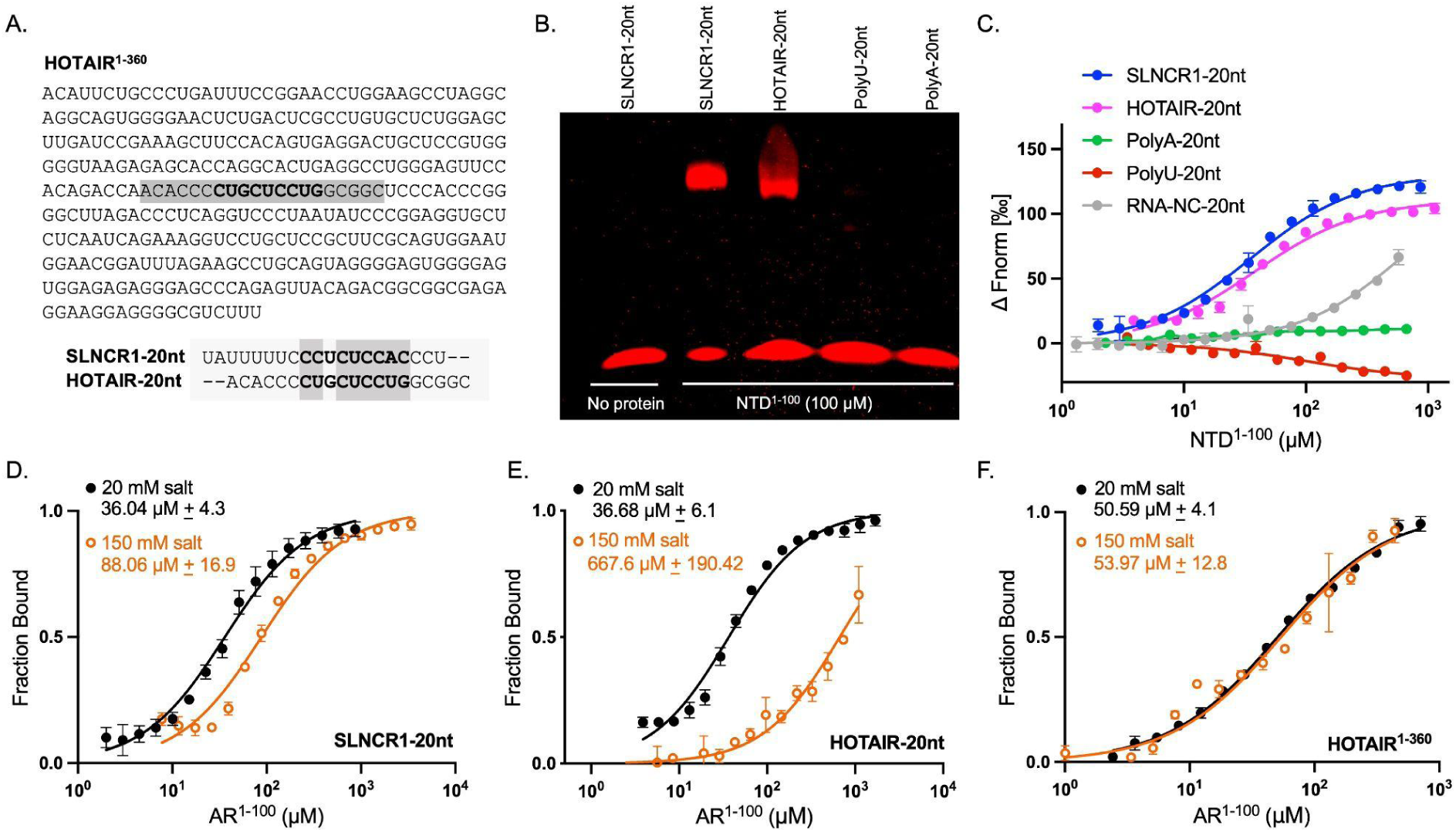
AR^1-100^ binds lncRNA fragments in a sequence-specific and partially electrostatic manner. A) Sequence of HOTAIR^1-360^ variant, with the sequence corresponding to the HOTAIR-20nt probe highlighted in gray. B) REMSA analysis of 100 μM AR^1-100^ binding to 50 nM fluorescently labeled 20-nucleotide RNA fragments resolved on a 6% TB gel. C) MST dose-response curve measurements for AR^1–100^ interaction with Cy5-labeled 20-nucleotide RNA oligonucleotides, including SLNCR1-20nt (blue), HOTAIR-20nt (magenta), PolyA-20nt (green), PolyU-20nt (red), and RNA-NC-20nt (gray). Measurements were performed under identical buffer and instrument conditions in 3 technical replicates. D, E, F) MST binding affinity measurements of AR^1-100^ interaction with Cy5-labeled SLNCR1-20nt (20 nM), HOTAIR-20nt (20 nM), and HOTAIR^1-360^ (50 nM) under different salt concentrations in 3 biological replicates, with corresponding Kd values provided.

To assess the extent to which the binding between SLNCR1-20nt or HOTAIR-20nt and AR^1-100^ is driven by electrostatics, we measured their salt dependence. Increasing the ionic strength from low salt (20 mM KCl) to near-physiological conditions (20 mM KCl + 130 mM NaCl) results in only a modest decrease in the binding affinity of AR^1-100^ for the SLNCR1-20nt RNA fragment (**Figure 3D**). In contrast, the binding of HOTAIR-20nt to AR^1-100^ weakens by an order of magnitude (but does not disappear) at higher ionic strength (**Figure 3E**). This divergent salt response suggests that SLNCR1-20nt binding is stabilized by additional non-electrostatic contacts, whereas the HOTAIR-20nt interaction relies more heavily on electrostatic contributions. Interestingly, this effect disappears when using the longer HOTAIR^1-360^, the binding of which seems insensitive to the increased ionic strength (**Figure 3F**). Thus, additional non-electrostatic contacts may occur between AR^1-100^ and the longer HOTAIR^1-360^ RNA, although they do not result in an intrinsically higher affinity.

To further examine whether the more pronounced salt dependence is due to the greater contribution of electrostatic forces to the interaction between AR^1-100^ and HOTAIR-20nt compared with SLNCR1-20nt, we performed molecular dynamics (MD) simulations of AR^1-100^ in the presence of both RNA fragments and a PolyA-20nt negative control. Starting from spatially separated random configurations, we analyzed interaction energy as a function of peptide-RNA distance during the first 1000 ns of the simulations. In both complexes, favorable Coulomb interaction energies are already observed at large distances, consistent with long-range electrostatic steering during the initial encounter **(Figure 4A, C)**. For the interaction between AR and HOTAIR-20nt, favorable Coulombic contributions prevail and strengthen as the interaction evolves toward shorter distances, indicating that recognition and early complex stabilization are dominated by electrostatic contributions **(Figure 4A, B)**. To avoid overinterpreting force-field-sensitive energetic terms, MD-derived Coulombic and Lennard-Jones energies were analyzed qualitatively as relative descriptors of RNA recognition and stabilization, rather than as quantitative estimates of binding affinity. Although the complex between AR and SLNCR1-20nt follows a similar electrostatically guided initial approach **(Figure 4C)**, Coulomb stabilization does not deepen to the same extent upon binding, and a larger contribution from Lennard-Jones interactions is also seen, consistent with a greater role of short-range non-electrostatic contacts in its stabilization **(Figure 4D)**. Consistent with its lack of detectable experimental binding, the PolyA-20nt forms no persistent contacts and only transiently approaches AR^1-100^ **(Figure 5C)**.

**Figure 4.**
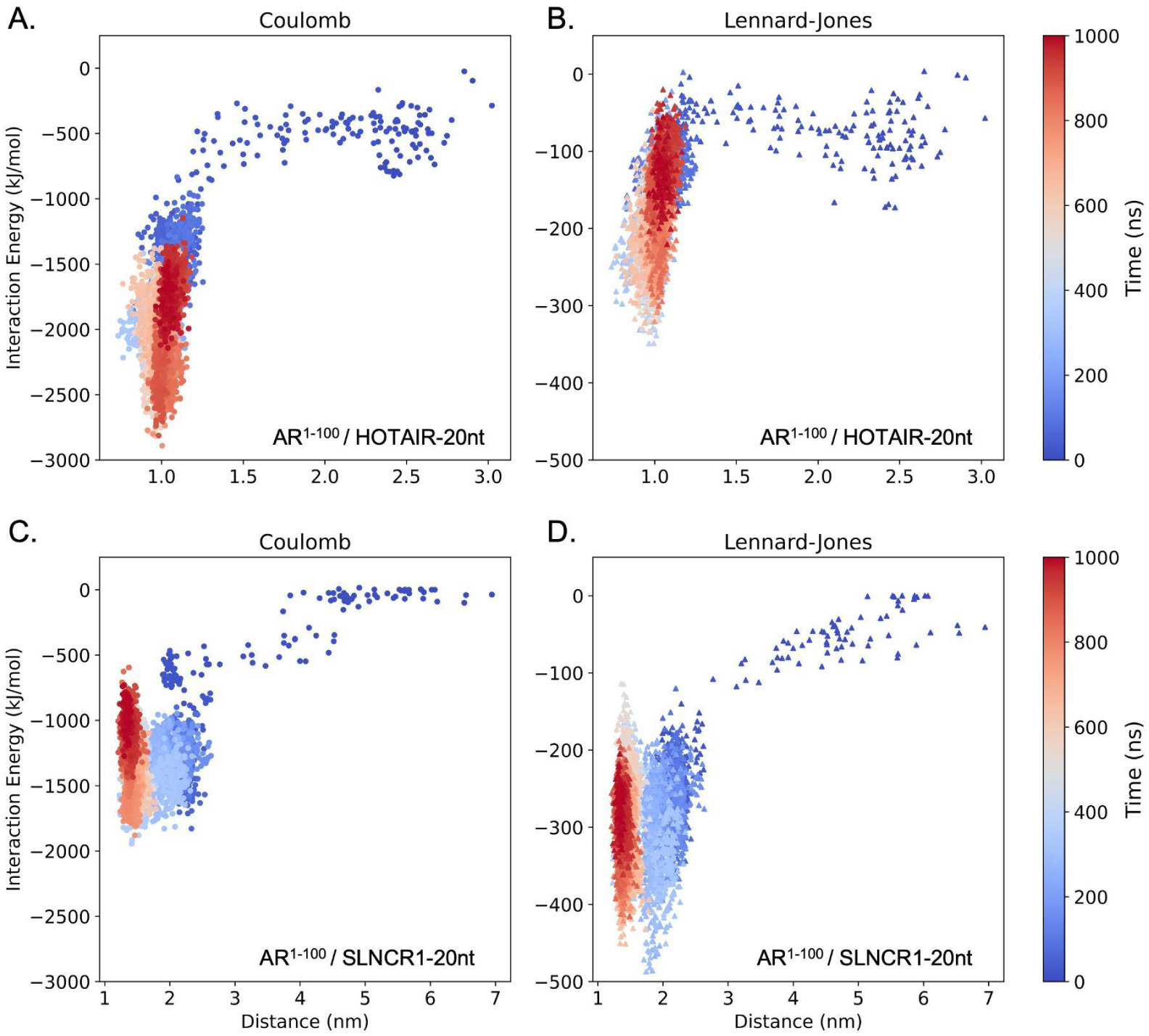
Distance-resolved interaction energy profiles of AR^1–100^ with HOTAIR-20nt and SLNCR1-20nt. A, B) Coulomb and Lennard-Jones interaction energy profiles of AR^1-100^ in complex with HOTAIR-20nt. C, D) Coulomb and Lennard-Jones interaction energy profiles of AR^1-100^ in complex with SLNCR1-20nt. Interaction energies are plotted as a function of the peptide-RNA epitope distance. points are colored by simulation time, illustrating the transition from the initial encounter to early-bound conformations.

**Figure 5.**
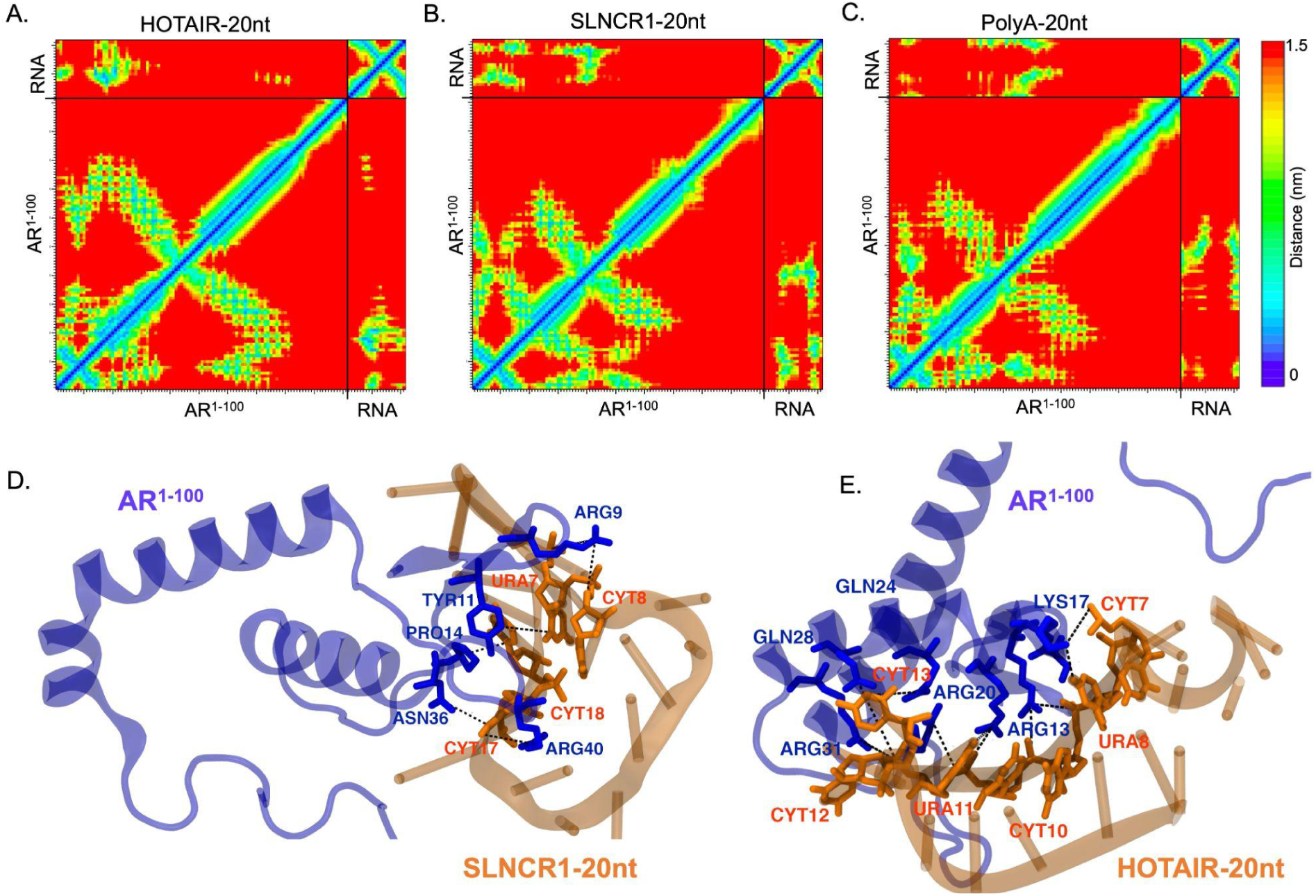
MD-based mapping of AR^NTD^ interactions with RNA fragments. A-C) Mean minimum distance contact maps for AR^1–100^ in complex with RNA fragments. Each heatmap reports, for every peptide-residue pair, the mean of the minimum peptide-RNA distances computed over the last 500 ns of the trajectory; lower values (blue) indicate closer approach, and higher values (yellow to red) indicate greater separation. The color scale spans 0-1.5 nm (see vertical bar). D, E) Representative snapshots of AR^1-100^ in complex with the corresponding RNA fragments, with the main contact sites highlighted.

To characterize these interaction modes at residue-level resolution, recurrent non-covalent contacts were analyzed using *RING* (47). Contacts above the 20% occupancy threshold support distinct interaction profiles for the two RNA fragments in both unbiased MD simulations and HADDOCK refined trajectories. For HOTAIR-20nt, the observed interaction network is relatively compact and mainly composed of van der Waals contacts, with additional cation-π interactions and hydrogen-bonding contributions. In contrast, the interaction of AR with SLNCR1-20nt displays a broader and more chemically diverse interaction pattern, including van der Waals contacts, hydrogen bonds, cation-π interactions, and π-stacking-like interactions. This broader repertoire of short-range interactions is consistent with the weaker salt dependence observed experimentally for SLNCR1-20nt compared with HOTAIR-20nt. Detailed residue-nucleotide pairs, interaction types, and occupancies derived from the unbiased and HADDOCK-refined trajectories are provided in **Supplementary Tables S4 and S5.**

### Mapping of the RNA-binding residues of AR^1-100^

The MD-derived mean minimum-distance contact maps identified the N-terminal region of AR^1-100^ as the principal RNA-binding surface, consistent with the independent HADDOCK docking calculations **(Figure 5A-C; Supplementary Figure S3)**. In unbiased MD simulations, HOTAIR-20nt forms a compact and persistent interaction region centered on residues 12-31, with recurrent contacts involving Arg13, Ser16, Lys17, Arg20, Gln24, Gln28, and Arg31 **(Figure 5A; Supplementary Table S4)**. In contrast, SLNCR1-20nt shows a broader and more heterogeneous binding ensemble, with contacts distributed across residues 9-18, including Arg9, Tyr11, Arg13, and Pro14, and a second region around Arg40 involving Gln35, Asn36, Pro39, and Arg40 **(Figure 5B; Supplementary Table S4)**. At the RNA level, the main AR-contacting region in HOTAIR-20nt involved nucleotides 6-13, clustered within the CUGCUCCUG motif, whereas SLNCR1-20nt contacted AR mainly through nucleotides 7-8 and 15-18 overlapping with its motif-containing region. Consistently, representative snapshots illustrate the interaction of AR^1-100^ with these AR-interacting nucleotides within the motif containing RNA fragments **(Figure 5D, E; Supplementary Figure S4)**.

MD refinements of the top-ranked HADDOCK poses provide complementary support for docking-compatible AR-RNA interfaces and confirmed recurrent involvement of the same AR N-terminal RNA-binding surface for both RNAs, while the main RNA-contacting nucleotides were partially shifted relative to the unbiased simulations **(Supplementary Figure S3; Supplementary Table S5)**. These findings support a dynamic RNA-IDR binding mode in which AR engages motif-containing RNA regions through recurrent protein hotspots, rather than a single fixed residue-nucleotide arrangement.

To underpin the *in silico* results, NMR spectroscopy was used to map the protein-RNA interaction at residue-level resolution via chemical-shift perturbation analysis. Two-dimensional [¹H–^15^N] HSQC spectra were recorded for the uniformly [¹³C,^15^N]-labeled AR^1-100^ in the absence and presence of increasing concentrations of SLNCR1-20nt or HOTAIR-20nt RNA fragments. Upon titration, gradual and residue-specific chemical shift perturbations were observed. Mapping these chemical shift perturbations onto the AR^1-100^ sequence reveals that the most pronounced shifts are localized in the segment R^9^VYPRPPSKTYRGAFQNLF^27^, which can therefore be considered as the primary binding region to both RNA fragments (**Figure 6A-D**). The interactions are in fast exchange on the NMR chemical shift timescale, as indicated by gradual, continuous chemical shift changes for F27, R9, R13, and Q24 upon RNA titration, consistent with a transient, dynamic RNA-protein binding (**Figure 6A, B**) and in agreement with the MD simulations **(Figure 5)**. The dissociation constants obtained by simultaneously fitting the changes in backbone amide chemical shifts of R9, Y11, Q24, and F27 are 35 ± 5 µM for SLNCR1-20nt and 88 ± 5 µM for HOTAIR-20nt, respectively (**Figure 6E, F**), slightly lower than the ones measured using MST.

**Figure 6.**
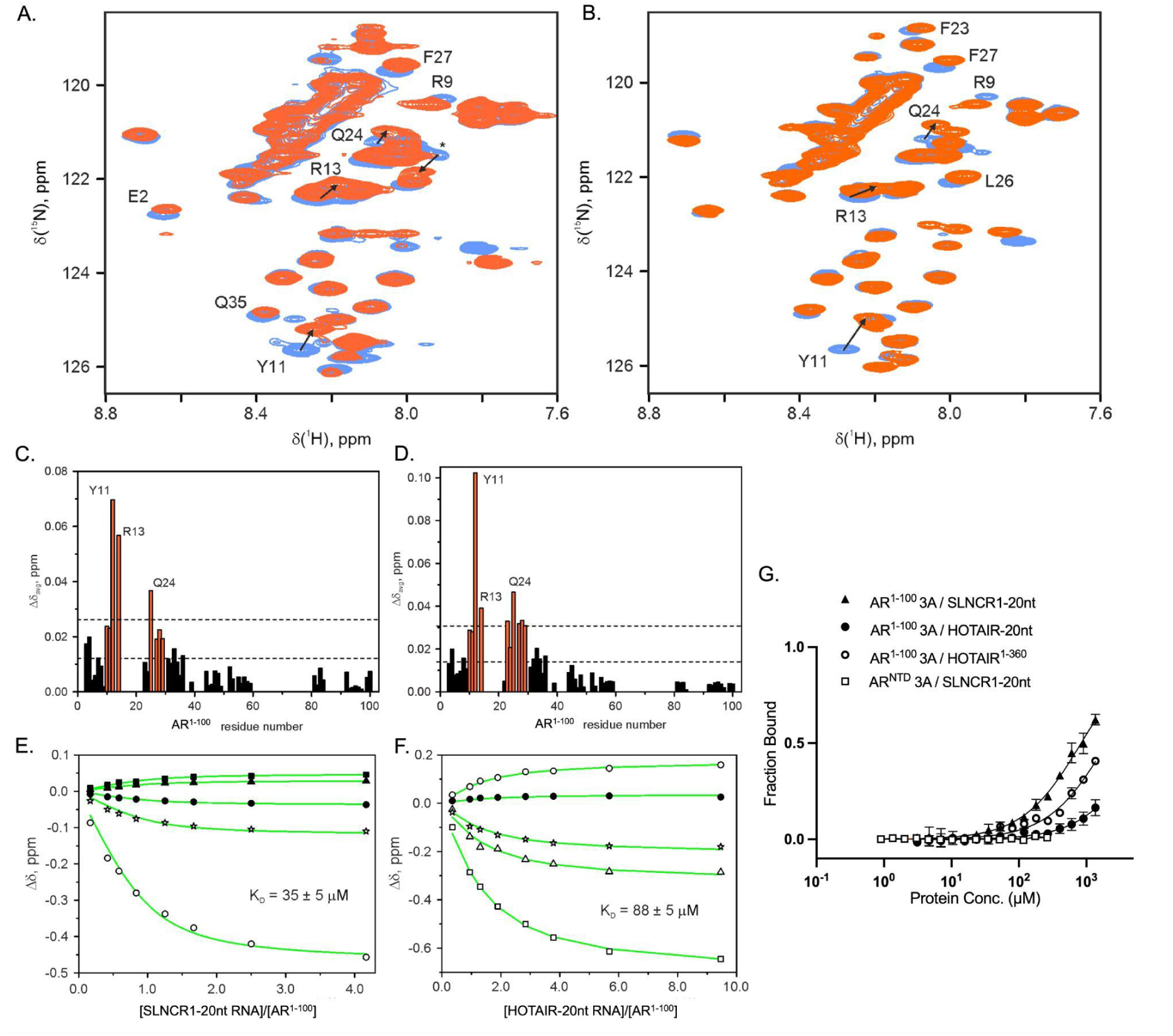
Mapping the minimal region for lncRNA binding within AR^1-100^. A, B) Chemical shift perturbations of AR^1-100^ backbone amide resonances upon binding to (A) SLNCR1-20nt and (B) HOTAIR-20nt. Free protein is shown in blue, while final titration points with 4.2 molar equivalents of SLNCR1-20nt and 5 molar equivalents of HOTAIR-20nt are shown in orange. The labels indicate the largest binding shifts; the asterisk denotes an unassigned resonance. C, D) Average binding shifts of AR^1-100^ backbone amides (Δδavg) in the presence of (C) 4.2 molar equivalents of SLNCR1-20nt or (D) 5 molar equivalents of HOTAIR-20nt RNA. The horizontal lines show the average Δδavg and the average plus one standard deviation. Protein regions with binding effects encompassing contiguous stretches of four or more residues are colored orange. The labels indicate residues with the largest Δδavg. E) Chemical shift perturbations (Δδ) of H (filled symbols) and N (open symbols) atoms of AR^1-100^ backbone amides Y11 (circles), R9 (triangles), V3 (stars), and unassigned resonance (labelled by an asterisk in Fig. 6A, squares) upon titration with SLNCR1-20nt were fitted simultaneously to a 1:1 binding model with the shared Kd (Eq. 1). The solid lines show the best fit with the Kd value of 35.1 ± 5.3 µM. F) Chemical shift perturbations (Δδ) of H (filled symbols) and N (open symbols) atoms of AR^1-100^ backbone amides R9 (circles), Y11 (squares), Q24 (triangles), and F27 (stars) upon titration with HOTAIR-20nt were fitted simultaneously to a 1:1 binding model with the shared Kd (Eq. 1). The solid lines show the best fit with the Kd value of 88.2 ± 5.3 µM. G) Binding of AR^1-100^-3A and AR^NTD^-3A to the indicated Cy5-labeled RNA fragments (20nM SLNCR1-20nt, 20 nM HOTAIR-20nt, and 50 nM HOTAIR^1-360^) was assessed by MST. Error bars represent the SD across three measurements, including three biological replicates.

Further validation comes from mutating Y11, R13, and Q24 to alanine (AR^1-100^-3A). This mutant shows markedly reduced binding to all tested RNA fragments, for SLNCR1-20nt the apparent Kd increases from 36.04 ± 4.39 μM to 833.84 ± 91.63 μM. Binding to HOTAIR-20nt and the longer HOTAIR^1-360^ fragment was strongly impaired and too weak to be reliably quantified, corresponding to lower-limit estimates in the millimolar range (Kd > 5 mM and Kd > 1 mM, respectively; **Figure 6G**). Introducing the same mutations in the full-length N-terminal domain (AR^NTD^-3A) equally abolishes binding to SLNCR1-20nt **(Figure 6G)**. Thus, these three amino acids form the core of the RNA-binding site on AR^1-100.^

### RNA-mediated phase separation of AR requires specific protein-RNA interactions

The intrinsically disordered AR^NTD^ plays a key role in LLPS of both the full-length protein and the AR-V7 isoform (18,20), the latter being linked to the pathomechanism of castration-resistant prostate cancer (23). As AR-interacting lncRNAs, including HOTAIR and SLNCR1, are upregulated in castration-resistant prostate cancer cells following androgen deprivation therapy, we tested whether their binding to AR^NTD^ can modulate its LLPS. We examined *in vitro* droplet formation using equimolar AR^NTD^ and Cy5-HOTAIR^1-360^ in the absence of macromolecular crowding agents, under biologically relevant buffer conditions (20 mM NaPi, pH 7.2, 200 mM NaCl, 1 mM MgCl₂, 0.5 mM TCEP). Under these conditions, neither AR^NTD^ nor Cy5-HOTAIR^1-360^ alone forms droplets. However, mixing AR^NTD^ with Cy5-HOTAIR^1-360^ triggers robust droplet formation, and the Cy5 signal strongly co-localizes with the resulting condensates **(Figure 7A)**. FRAP analysis of AR^NTD^ and HOTAIR condensates shows rapid fluorescence recovery, consistent with their liquid-like nature **(Figure 7B)**. In contrast to HOTAIR^1-360^, Cy5-labeled PolyU, HOTAIR-20nt, and SLNCR1-20nt RNA do not promote droplet formation under identical conditions **(Figure 7A)**. Moreover, the AR^NTD^ mutant carrying Y11A, R13A, and Q24A mutations (AR^NTD^-3A) fails to form condensates in the presence of HOTAIR^1-360^ **(Figure 7C)**. Together, these results show that HOTAIR lncRNA promotes AR^NTD^ LLPS through a sequence-dependent mechanism, rather than through nonspecific electrostatic effects of RNA.

**Figure 7.**
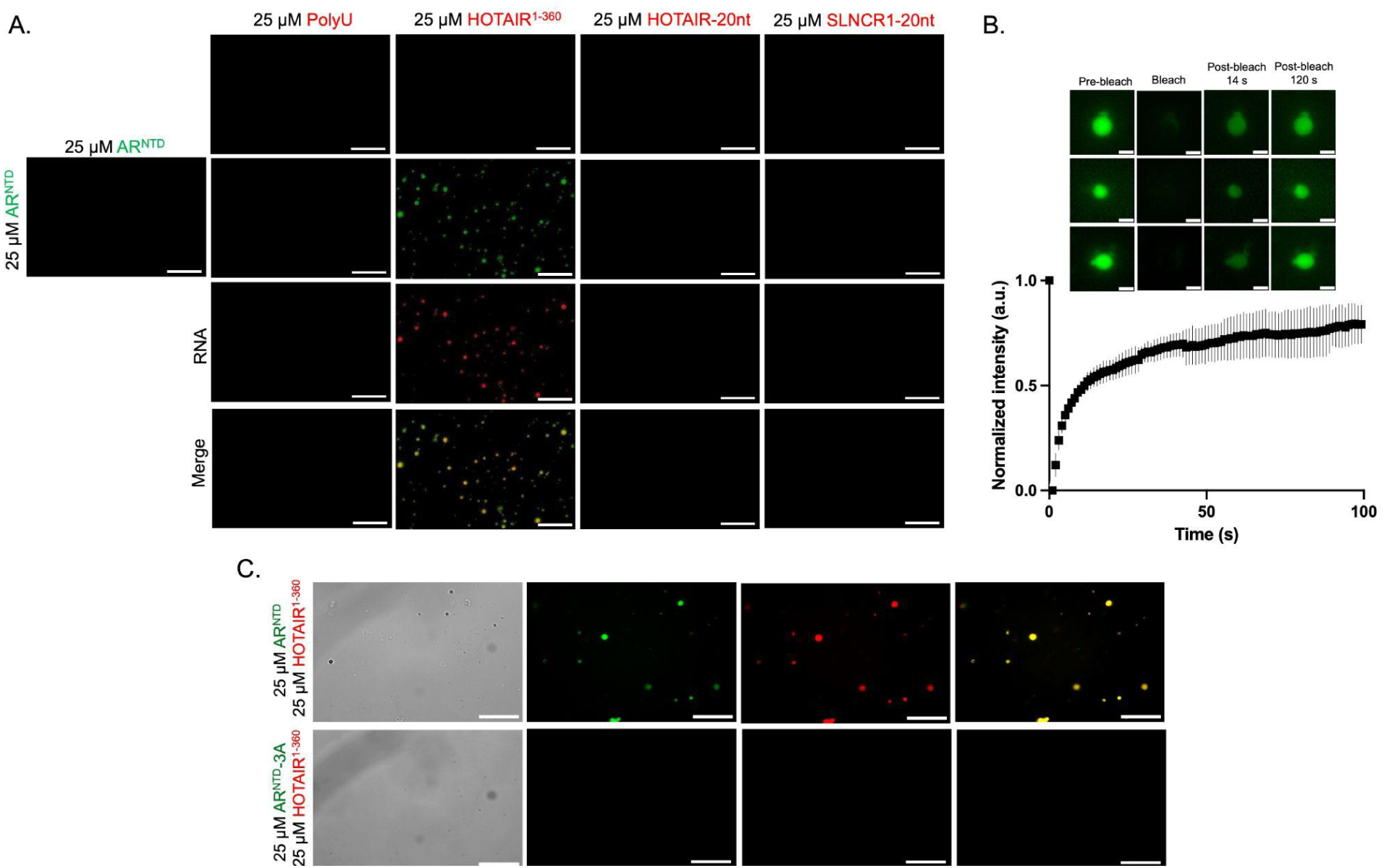
RNA-dependent phase behavior of AR^NTD^. A) Fluorescence microscopy images of 25 μM DyLight488-labeled AR^NTD^ in the presence of 25 μM Cy5-labeled PolyU or HOTAIR^1-360^ or HOTAIR-20nt or SLNCR1-20nt. Droplet formation is observed in the presence of HOTAIR^1-360^, which colocalizes with AR^NTD^ (scale bar = 10 μm; similar results observed in three biological replicates). B) FRAP analysis of droplets formed by 25 μM AR^NTD^ and 25 μM HOTAIR^1–360^ (scale bar= 1 μm, 4 technical replicates). C) Fluorescence microscopy images of 25 μM DyLight488-labeled AR^NTD^ and 25 μM AR^NTD^ 3A mutant in the presence of 25 μM HOTAIR^1-360^ (scale bar = 10 μm; similar results observed in three biological replicates). All LLPS experiments were performed in 20 mM NaPi, pH 7.2, 200 mM NaCl, 1 mM MgCl₂, and 0.5 mM TCEP.

### LncRNA and AR^LBD^ compete for binding to AR^NTD^

In the context of full-length AR, residues F^23^QNLF^27^ of the NTD form a linear motif that docks into the AF-2 groove of the ligand-binding domain when the latter is in its DHT- or testosterone-bound state (48). As this motif partially overlaps with the above identified lncRNA-binding region of the NTD, we investigated whether the two interactions may compete or reinforce each other. We first compared the binding of AR^1-100^ and the AR^1-100^ 3A mutant (carrying the Y11A, R13A, and Q24A mutations) to the AR^LBD^ in the absence of RNA. The 3A mutant binds the AR^LBD^ with nearly a tenfold lower affinity, with a Kd of 164.41 μM ± 20.32, compared to 17.79 μM ± 1.44 for the wild-type protein AR^1-100^ **(Figure 8A)**, in agreement with this region being important for N/C interdomain communication. Consistent with this, the AR^NTD^ 3A mutant forms much smaller condensates in the presence of AR^LBD^ than the wild-type AR^NTD^ **(Figure 8B)**. FRAP analysis further confirmed the dynamic behavior of wild-type condensates **(Supplementary Figure S2)**, in line with previous evidence that N/C interdomain coupling supports AR phase separation (20,22).

**Figure 8.**
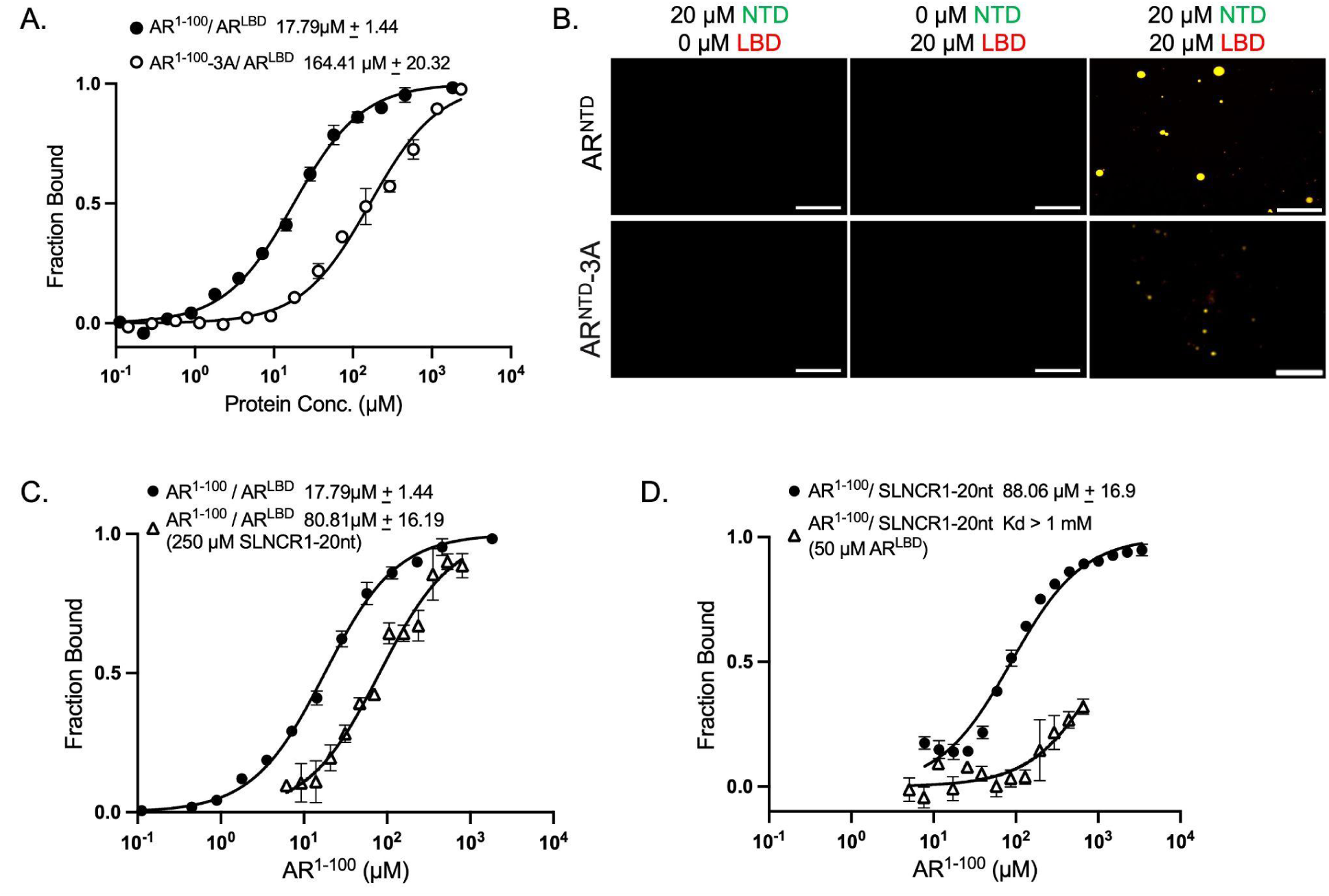
The NMR-mapped RNA-binding residues are implicated in AR N/C interdomain communication. A) MST binding affinity measurements of AR^1-100^ and AR^1-100^ 3A interaction with Cy5-labeled AR^LBD^ (33nM). Error bars indicate SD of triplicate measurements (two biological and one technical replicates). B) Fluorescence microscopy images of purified full-length AR^NTD^ constructs (WT and 3A mutant), the AR^LBD^, and an equimolar mixture of both proteins (20 μM). The green and red channels, corresponding to AR^NTD^ and AR^LBD^, respectively, are shown as merged images. (scale bar = 10 μm; similar results observed in three biological replicates). C) MST binding affinity measurements of AR^1-100^-AR^LBD^ binding in the presence and absence of 250 μM SLNCR1-20nt, and D) AR^1-100^-SLNCR1-20nt binding in the presence and absence of 50 μM AR^LBD^. Error bars indicate SD of three biological replicates. Assay buffer: 20 mM NaPi, pH 7.2, 200 mM NaCl, 0.5 mM TCEP, and 10 μM DHT.

**Figure 9.**
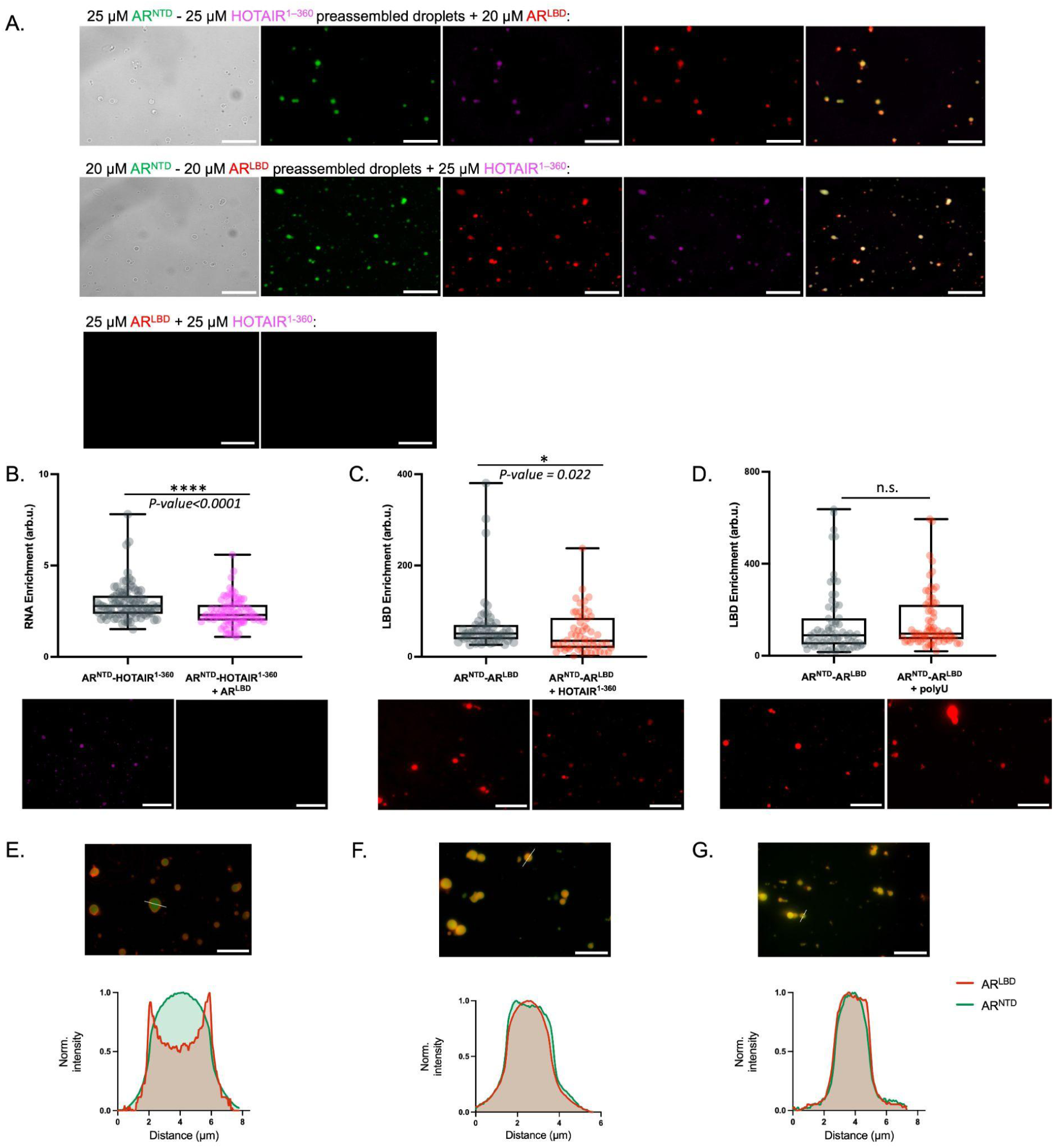
LncRNA-mediated modulation of AR N/C interdomain communication. A) Fluorescence microscopy images of preassembled equimolar condensates showing reciprocal recruitment of HOTAIR^1-360^ and AR^LBD^. Upper: DyLight488-labeled AR^NTD^ and Cy5-labeled AR^LBD^ droplets after addition of 25 μM Cy3-labeled HOTAIR^1-360^. Lower: DyLight488-labeled AR^NTD^ and Cy3-labeled HOTAIR^1-360^ droplets after addition of 20 μM Cy5-labeled AR^LBD^. An equimolar mixture of Cy5-labeled AR^LBD^ and Cy3-labeled HOTAIR^1-360^ (25 μM each) did not undergo LLPS. Scale bar, 10 μm. Representative of three biological replicates. B) Quantification of Cy3-labeled HOTAIR^1-360^ enrichment in preassembled AR^NTD^-HOTAIR^1-360^ droplets before and after the addition of 20 μM Cy5-labeled AR^LBD^. C) Quantification of Cy5-labeled AR^LBD^ enrichment in preassembled AR^NTD^-AR^LBD^ droplets before and after the addition of 25 μM Cy3-labeled HOTAIR^1-360^. D) Quantification of Cy5-labeled AR^LBD^ enrichment in preassembled AR^NTD^-AR^LBD^ droplets before and after the addition of an equivalent amount of PolyU-20nt. For B-D, enrichment was calculated from raw-image fluorescence measurements. Boxes indicate quartiles and the mean; dots represent individual droplets across three biological replicates. Representative images are shown with identical linear scaling within each channel across conditions. E-G) Fluorescence microscopy images from three biological replicates of preassembled equimolar DyLight488-labeled AR^NTD^ and Cy5-labeled AR^LBD^ droplets after the addition of E) 400 μM SLNCR1-20nt, F) HOTAIR-20nt, or G) PolyU-20nt. Scale bar, 10 μm. The corresponding normalized line-scan intensity profiles are shown.

We next probed the interaction between AR^1-100^ and AR^LBD^ in the absence and presence of SLNCR1-20nt. The observed affinity of AR^1-100^ for AR^LBD^ decreases approximately 5-fold in the presence of 250 μM SLNCR1-20nt, from 17.79 ± 1.44 μM to 80.81 ± 16.19 μM **(Figure 8C)**, indicating competition for the same or overlapping binding sites. This is further confirmed by the inverse experiment, in which the presence of 50 μM AR^LBD^ strongly perturbs the AR^1-100^-SLNCR1-20nt interaction **(Figure 8D)**. Together, these results suggest mutual interference between SLNCR1-20nt and AR^LBD^ binding to the NTD.

### LncRNA binding affects AR N/C interdomain association within condensates

We next asked whether lncRNA and AR^LBD^ can coexist in AR^NTD^ droplets or whether one would exclude the other. Upon addition of 25 μM AR^LBD^ to preassembled equimolar AR^NTD^ and HOTAIR^1-360^ droplets (25 μM each), AR^LBD^ incorporates into the condensates **(Figure 9A, top panel)**. At the same time, RNA enrichment in the condensates becomes significantly reduced, mirroring the competition between lncRNA and AR^LBD^ for binding to NTD **(Figure 9B)**. Reciprocally, addition of 25 μM Cy3-labeled HOTAIR^1-360^ to preformed equimolar AR^NTD^ and AR^LBD^ droplets (20 μM each) results in its colocalization with both AR^NTD^ and AR^LBD^ **(Figure 9A, lower panel)** and reduces the AR^LBD^ signal within the condensates **(Figure 9C)**. In contrast, the addition of nonspecific PolyU RNA at a nucleotide-equivalent amount to 25 μM HOTAIR^1-360^ does not produce a similar effect in AR^NTD^-AR^LBD^ droplets **(Figure 9D),** confirming that the specific interaction between lncRNA and AR^NTD^ is essential and that they compete with each other for interaction with AR^NTD^ within condensates. If insufficient AR^NTD^ binding sites are available, the non-specific multivalent interactions needed to stabilize condensates are insufficient to keep the excess lncRNA or AR^LBD^ in the droplets.

To further confirm that these effects are due to the specific binding of the C[CU]U[CU]UCC[AU][GC] motif to AR^NTD^ rather than non-specific interactions with the RNA, we carried out a similar experiment in which 400 μM SLNCR1-20nt, HOTAIR-20nt, or PolyU-20nt was added to preassembled equimolar AR^NTD^-AR^LBD^ droplets (20 μM each). SLNCR1-20nt addition induces a halo-like redistribution of AR^LBD^, with Cy5-AR^LBD^ appearing enriched at the droplet periphery rather than uniformly distributed throughout the condensate **(Figure 9E).** By contrast, HOTAIR-20nt does not detectably redistribute AR^LBD^ under these conditions **(Figure 9F)**, likely reflecting the salt sensitivity of the HOTAIR-20nt interaction in the LLPS buffer containing 200 mM NaCl. Similarly, PolyU-20nt had no detectable effect on AR^LBD^ distribution **(Figure 9G)**. Together, these results indicate that lncRNAs can coexist with AR^LBD^ in AR^NTD^ condensates but alter condensate composition and internal organization in an RNA-specific manner. In particular, SLNCR1-20nt remodels AR^NTD^-AR^LBD^ condensates without fully excluding AR^LBD^ from the condensed phase. Thus, RNA binding competes with the AR N/C interdomain association, consistent with competition for overlapping binding regions in solution.

## Discussion

The interaction between the androgen receptor and long non-coding RNAs plays an important role in prostate cancer. It enhances AR transcriptional activity toward target genes, drives tumor growth, and modulates AR signaling by influencing its localization, co-factor recruitment, chromatin association, or receptor stability (13,15–17). However, how AR recognizes lncRNAs in a sequence-specific manner and the mechanism by which this interaction communicates to the ligand-binding domain remains poorly understood. Our combined NMR and *in silico* investigations, together with biophysical assays, suggest that motif-containing regions of HOTAIR and SLNCR1 lncRNAs specifically interact with the N-terminal domain of AR, with the primary contact points being between residues 10 and 30 of AR^NTD^. Although the interaction is specific on both the RNA and protein sides, it is a "fuzzy" interaction in which both partners remain fully disordered. Using NMR chemical-shift analysis and site-specific mutagenesis, we identified residues Y11, R13, and Q24 as key determinants of RNA recognition. Computational analyses further support this model, with both unbiased MD simulations and HADDOCK-derived refinements identifying the experimentally defined AR^NTD^ binding hotspot and suggesting dynamic, partially variable RNA contacts consistent with fuzzy RNA-IDR recognition.

The short patch of lncRNA-interacting residues overlaps with the well-described F^23^QNLF^27^ motif, which mediates crucial interdomain communication with the ligand-binding domain of AR (33), suggesting that AR-interacting lncRNAs play a regulatory role in interdomain communication and coactivator recruitment. The F^23^QNLF^27^ motif engages the AF-2 surface of AR^LBD^ (residues 665-920) by adopting an α-helical conformation upon binding, representing a folding-upon-binding mechanism (49,50) that is fundamentally different from the fuzzy interaction with lncRNAs observed through NMR. Such a partner-dependent change in binding mode is rare, with only a few examples described in the literature. One of the strongest cases is the IDR of the *Vibrio cholerae* antitoxin HigA2, which folds upon binding to its cognate toxin HigB2 but also interacts in a fully fuzzy manner with the operator DNA (51). Experimental evidence for a similar behavior is available for at least three other proteins, although structural characterization is more limited. A segment within the measles virus NTAIL folds into an a-helix upon binding the X domain of the viral phosphoprotein, while likely remaining disordered when interacting with hsp70 (52). Similarly, the intrinsically disordered WW domain of KIBRA folds into the canonical β-sheet with Dendrin, but remains largely disordered when bound to synaptopodin (53). Finally, the intermediate chain subunit of Dynein shows increasing structure upon binding p150*Glued,* while order decreases upon binding NudE (54). Other cases correspond to IDRs that adopt different ensembles upon (partially) folding upon binding to distinct partners, but do not remain fully disordered.

Our mutation and MST assays show that the NMR-defined RNA-binding residues contribute to AR^1-100^ and AR^LBD^ association, and that SLNCR1-20nt and AR^LBD^ mutually interfere with binding to AR^1-100^, consistent with partially overlapping interaction surfaces. In this reconstituted NTD-LBD system, however, the apparent Kd values and RNA-mediated competition should not be interpreted as quantitative measures of cis N/C communication in full-length. Instead, these data indicate that lncRNA binding can modulate LBD engagement through an overlapping interaction region within the AR^NTD^. Thus, lncRNAs upregulated in therapy-adapted prostate cancer may influence AR interdomain coupling by engaging the AR^NTD^ region that partially overlaps with the F^23^QNLF^27^ motif, potentially affecting AF-2 accessibility and coactivator recruitment. These effects may be particularly relevant under low-androgen conditions, where ligand-dependent N/C association is weakened or dynamically regulated, as is the case with the castration-resistant prostate cancer-associated AR coregulator MAGEA11, which regulates AR allostery under low-androgen conditions (55,56).

In AR variants such as AR-V7, which lack the LBD, lncRNA binding would not affect canonical N/C communication directly, but may instead affect NTD-mediated regulatory mechanisms, including condensate formation and ligand-independent transcriptional activity. In this context, the effect of AR-lncRNA interactions on AR phase separation becomes particularly important. The intrinsically disordered AR^NTD^ is a major contributor to phase separation of both full-length AR and the truncated AR-V7 isoform (18,20). We show that HOTAIR^1-360^, which contains the AR-interacting motif, specifically promotes LLPS of the AR^NTD^ in the absence of any crowding agent. These findings suggest that specific lncRNAs can lower the LLPS threshold of the AR^NTD^ and may therefore contribute to context-dependent AR condensate formation in therapy-adapted prostate cancer states. This may be specifically relevant to AR-V7 condensate-dependent activation of oncogenic transcriptional programs in castration-resistant prostate cancer, as AR-V7 has a higher concentration threshold for condensate formation than full-length AR, both in cells and in vitro, and its LLPS capacity is strongly shaped by cellular context (23).

Full-length AR is known to form nuclear condensates upon androgen stimulation in androgen-sensitive prostate cancer cells (18,20,22). In our reconstituted AR^NTD^-AR^LBD^ system, HOTAIR^1-360^ colocalizes with AR^NTD^-AR^LBD^ condensates while reducing, but not abolishing, AR-LBD enrichment. Together, these results suggest that lncRNAs can remodel condensate composition by introducing additional multivalent RNA-mediated interaction surfaces and tuning N/C interdomain association.

Our findings fit within a broader model in which androgen deprivation therapy exerts opposing effects on AR activity. Although androgen deprivation therapy directly suppresses androgen-dependent AR transactivation, long-term therapy adaptation can indirectly boost AR signaling through mechanisms such as the emergence of androgen-independent AR isoforms (5) and upregulation of androgen-repressed lncRNAs, such as HOTAIR, LINC00675, PCAT1, SOCS2-AS1, PCGEM1, and PRNCR1 (12–17). This suggests that hormone therapy should be complemented by strategies that counteract these adaptive mechanisms to ultimately prevent the development of castration-resistant prostate cancer (57–60). Selective AR-irreversible covalent antagonists (SARICAs), which target the AR transactivation domain within the AR^NTD^, represent one such approach (61). In this context, our finding that SLNCR1 and HOTAIR converge on the same RNA-binding surface within the AR^NTD^ highlights this region as a potential targetable interface for modulating AR-lncRNA interactions, which may complement approaches aimed at individual lncRNAs, such as antisense targeting of SLNCR1 (43). Together, our in vitro system defines a direct molecular mechanism linking AR-lncRNA recognition to RNA-dependent condensate remodeling, providing a foundation for future cellular and in vivo studies to determine how this mechanism functions within the native AR regulatory environment.

## Author contributions

**Nazanin Farahi:** Conceptualization, investigation, Formal analysis, Funding acquisition, Methodology, Visualization, Writing—original draft, Writing—review & editing; **Galo Ezequiel Balatti:** investigation; Formal analysis, Methodology, Visualization, Writing—review & editing; **Alexander N. Volkov:** Investigation; Formal analysis, Methodology, Visualization, Writing—review & editing; **Rita Pancsa:** Investigation, Writing—review & editing**; Peter Tompa:** Conceptualization, Supervision, Project administration, Writing—original draft, Writing—review & editing**; Remy Loris:** Conceptualization, Supervision, Project administration, Writing—original draft, Writing—review & editing.

## Acknowledgements

The authors gratefully acknowledge Prof. Dr. Ulrich Hennecke from the Organic Chemistry Research Group (ORGC) at the Vrije Universiteit Brussel (VUB) for his assistance with HPLC analysis of the RNA samples, which was used to assess RNA purity and integrity.

## Data availability

The data underlying this paper are available in the article and its online supplementary file.

## Supplementary data

Supplementary data is available at NAR online.

