## Supplementary figures and tables for "Binding of long non-coding RNAs within the androgen receptor N-terminal tail modulates interdomain communication and liquid-liquid phase separation"

### Supplementary Tables

| **AR protein constructs** | **Amino acid sequence** |
|---|---|
| $AR^{1-100}$ | MEVQLGLGRVYPRPPSKTYRGAFQNLFQSVREVIQNPGPRHPEAASAAPPGASLLL LQQQQQQQQQQQQQQQQQQQQQQQETSPRQQQQQGEDGSPQAH |
| $AR^{NTD}$ | MEVQLGLGRVYPRPPSKTYRGAFQNLFQSVREVIQNPGPRHPEAASAAPPGASLLL LQQQQQQQQQQQQQQQQQQQQQQQETSPRQQQQQGEDGSPQAHRRGPT GYLVLDEEQQPSQPQSALECHPERGCVPEPGAAVAASKGLPQQLPAPPDEDDSAAP STLSLLGPTFPGLSSCSADLKDILSEASTMQLLQQQQQEAVSEGSSSGRAREASGAPT SSKDNYLGGTSTISDNAKELCKAVSVSMGLGVEALEHLSPGEQLRGDCMYAPLLGVP PAVRPTPCAPLAECKGSLLDDSAGKSTEDTAEYSPFKGGYTKGLEGESLGCSGSAAAG SSGTLELPSTLSLYKSGALDEAAAYQSRDYYNFPLALAGPPPPPPPPHPHARIKLENPL DYGSAWAAAAAQCRYGDLASLHGAGAAGPGSGSPSAAASSSWHTLFTAEEGQLYG PCGGGGGGGGGGGGGGGGGGGGGGGEAGAVAPYGYTRPPQGLAGQESDFTAP DVWYPGGMVSRVPYPSPTCVKSEMGPWMDSYSGPYGDMRLETARDHVLPIDYYF |
| $AR^{LBD}$ | EGYECQPIFLNVLEAIEPGVVCAGHDNNQPDSFAALLSSLNELGERQLVHVVKWAKA LPGFRNLHVDDQMAVIQYSWMGLMVFAMGWRSFTNVNSRMLYFAPDLVFNEYR MHKSRMYSQCVRMRHLSQEFGWLQITPQEFLCMKALLLFSIIPVDGLKNQKFFDEL RMNYIKELDRIIACKRKNPTSCSRRFYQLTKLLDSVQPIARELHQFTFDLLIKSHMVSV DFPEMMAEIISVQVPKILSGKVKPIYFHTQ |
| **RNA oligos** | **Sequence (5’....3’)** |

| | |
|---|---|
| SLNCR1-20nt | UAUUUUUCCCUCUCCACCCU |
| HOTAIR-20nt | ACACCCCUGCUCCUGGCGGC |
| RNA-NC-20nt | ACUCUCUCUCUCUCUAUCUC |
| PolyU-20nt | UUUUUUUUUUUUUUUUUUUU |
| PolyA-20nt | AAAAAAAAAAAAAAAAAAAA |
| HOTAIR$^{1-360}$ | ACAUUCUGCCCUGAUUUCCGGAACCUGGAAGCCUAGGCAGGCAGUGGGGAA CUCUGACUCGCCUGUGCUCUGGAGCUUGAUCCGAAAGCUUCCACAGUGAGG ACUGCUCCGUGGGGGUAAGAGAGCACCAGGCACUGAGGCCUGGGAGUUCCA CAGACCAACACCCCUGCUCCUGGCGGCUCCCACCCGGGACUUAGACCCUCAG GUCCCUAAUAUCCCGGAGGUGCUCUCAAUCAGAAAGGUCCUGCUCCGCUUC GCAGUGGAAUGGAACGGAUUUAGAAGCCUGCAGUAGGGGAGUGGGGAGUG GAGAGAGGGAGCCCAGAGUUACAGACGGCGGCGAGAGGAAGGAGGGGCGUC UUU |

**Supplementary Table S1.** Sequences of the protein fragments and RNA oligonucleotides used in this study.

| **Primer name** | **Primer sequence (5'....3')** |
|---|---|
| linear_Hotair360_F | ACCCACTGCTTACTGGCTTATCG |
| linear_Hotair360_R | TCCTCTCGCCGCCGTCTGTAAC |
| NTD1-100_R | ATGTGCCTGCGGGCTGCCGTCTTC |
| NTD1-100_F | TGACTGTGCGCGTCCCGTAATG |
| LBD_F | CACATCGAGGGTTACGAGTG |
| LBD_R | CATGCCTTGAAAGTACAGATTCTC |
| NTD1-100-Y11A_R | CCAAACCCAGCTGGACTTCC |
| NTD1-100-Y11A_F | GTCGTGTTGCTCCGAGACCACC |
| NTD1-100-R13A_F | GCTCCACCCAGCAAGACGTAC |
| NTD1-100-R13A_R | CGGAGCAACACGACCCAAAC |
| NTD1-100-Q24A_R | ACGCACCGCGGTACGTCTTGC |
| NTD1-100-Q24A_F | TTGCTAACCTGTTCCAGAGCGTTCGCGAGG |

**Supplementary Table S2. Primer sequences used for cloning and site-directed mutagenesis in this study.** All unmodified DNA oligonucleotides were designed in-house and purchased from Eurofins Genomics (Ebersberg, Germany). Primers were supplied lyophilized, resuspended in nuclease-free water, and diluted to 10 µM working stocks for PCR reactions.

| Query Motif | lncRNA | Strand | Start | End | p-value | q-value | Matched Seq |
|---|---|---|---|---|---|---|---|
| CYUCUCCWS | HOTAIR | + | 167 | 175 | 0.000671 | 0.233 | **CUGCUCCUG** |
| CYUCUCCWS | SLNCR1 | + | 45 | 53 | 3.05e-05 | 0.000931 | **CCUCUCCAC** |
| CYUCUCCWS | SLNCR1 | + | 60 | 68 | 3.05e-05 | 0.000931 | **CUUCUCCUG** |
| CYUCUCCWS | LINC00675 | + | 175 | 183 | 0.000671 | 0.161 | **UUUCUCCUC** |
| CYUCUCCWS | LINC00675 | + | 328 | 336 | 0.000671 | 0.161 | **UCUCUCCUC** |
| CYUCUCCWS | LINC00675 | + | 331 | 339 | 0.000671 | 0.161 | **CUCCUCCUC** |
| CYUCUCCWS | PCAT | + | 1670 | 1678 | 3.05e-05 | 0.0605 | **CUUCUCCAC** |
| CYUCUCCWS | PCAT | + | 174 | 182 | 0.000671 | 0.666 | CCUCUCUUG |
| CYUCUCCWS | SOCS2-AS1 | + | 307 | 315 | 6.1e-05 | 0.0326 | **CUUUUCCAC** |
| CYUCUCCWS | PRNCR | + | 7429 | 7437 | 3.05e-05 | 0.318 | **CUUCUCCUC** |
| CYUCUCCWS | PRNCR | + | 1346 | 1354 | 6.1e-05 | 0.318 | CUUUUCCUG |
| CYUCUCCWS | PRNCR | + | 1921 | 1929 | 0.000153 | 0.318 | CCUCUCCUU |
| CYUCUCCWS | PRNCR | + | 2794 | 2802 | 0.000153 | 0.318 | CAUCUCCAC |
| CYUCUCCWS | PRNCR | + | 8757 | 8765 | 0.000153 | 0.318 | CAUCUCCUG |
| CYUCUCCWS | PRNCR | + | 11200 | 11208 | 0.000153 | 0.318 | CCUCUCCAU |
| CYUCUCCWS | PRNCR | + | 1215 | 1223 | 0.000671 | 0.365 | CUUCGCCUC |
| CYUCUCCWS | PRNCR | + | 1359 | 1367 | 0.000671 | 0.365 | ACUCUCCAG |
| CYUCUCCWS | PRNCR | + | 1862 | 1870 | 0.000671 | 0.365 | UUUCUCCUG |
| CYUCUCCWS | PRNCR | + | 2479 | 2487 | 0.000671 | 0.365 | CCUCCCCUC |
| CYUCUCCWS | PRNCR | + | 2580 | 2588 | 0.000671 | 0.365 | UUUCUCCAG |
| CYUCUCCWS | PRNCR | + | 3419 | 3427 | 0.000671 | 0.365 | CUUCUACUG |
| CYUCUCCWS | PRNCR | + | 3602 | 3610 | 0.000671 | 0.365 | ACUCUCCAG |

| | | | | | | | |
|---|---|---|---|---|---|---|---|
| CYUCUCCWS | PRNCR | + | 5087 | 5095 | 0.000671 | 0.365 | CUUCUUCAG |
| CYUCUCCWS | PRNCR | + | 5542 | 5550 | 0.000671 | 0.365 | CCUCCCCAG |
| CYUCUCCWS | PRNCR | + | 5925 | 5933 | 0.000671 | 0.365 | CUUCUUCUG |
| CYUCUCCWS | PRNCR | + | 6057 | 6065 | 0.000671 | 0.365 | AUUCUCCAC |
| CYUCUCCWS | PRNCR | + | 8735 | 8743 | 0.000671 | 0.365 | CUUCUUCUC |
| CYUCUCCWS | PRNCR | + | 8738 | 8746 | 0.000671 | 0.365 | CUUCUCUUC |
| CYUCUCCWS | PRNCR | + | 8908 | 8916 | 0.000671 | 0.365 | CUUCUACUC |
| CYUCUCCWS | PRNCR | + | 9007 | 9015 | 0.000671 | 0.365 | CUUCUCAUC |
| CYUCUCCWS | PRNCR | + | 11467 | 11475 | 0.000671 | 0.365 | GUUCUCCUC |
| CYUCUCCWS | PRNCR | + | 12188 | 12196 | 0.000671 | 0.365 | CUACUCCAC |
| CYUCUCCWS | PRNCR | + | 6573 | 6581 | 0.000763 | 0.367 | CUUUUCCUU |
| CYUCUCCWS | PRNCR | + | 7088 | 7096 | 0.000763 | 0.367 | CCUUUCCUA |
| CYUCUCCWS | PRNCR | + | 9061 | 9069 | 0.000763 | 0.367 | CCUUUCCAU |
| CYUCUCCWS | PCGEM | + | 53 | 61 | 0.00287 | 0.155 | **CUACUCCUA** |

**Supplementary Table S3.** Candidate SLNCR1-like motifs were identified in AR-associated lncRNAs using FIMO from the MEME Suite v5.5.9 with the 9-nt consensus motif CYUCUCCWS as the query. The table lists the identified motif occurrences, including transcript name, strand, motif position, FIMO p-value and q-value, and matched sequence. Motifs prioritized for experimental follow-up, based on the lowest p-value hits, are highlighted in gray.

| AR | HOTAIR20 | Interaction | Occupancy (%) |
|---|---|---|---|
| ARG13 | CYT10 | vdW | 26.15 |
| ARG13 | GUA9 | vdW | 21.56 |
| SER16 | URA8 | vdW | 75.45 |
| SER16 | GUA9 | vdW | 29.34 |
| LYS17 | CYT7 | vdW | 45.31 |
| LYS17 | URA8 | H-BOND | 29.14 |
| LYS17 | URA8 | vdW | 26.15 |

| | | | |
|---|---|---|---|
| LYS17 | CYT6 | vdW | 22.16 |
| ARG20 | CYT10 | CATION-π | 69.46 |
| ARG20 | CYT10 | vdW | 42.91 |
| ARG20 | URA11 | vdW | 42.32 |
| GLN24 | CYT13 | vdW | 31.74 |
| GLN24 | CYT13 | H-BOND | 24.15 |
| GLN28 | CYT13 | vdW | 26.55 |
| ARG31 | CYT12 | vdW | 24.75 |
| ARG31 | CYT12 | CATION-π | 22.95 |
| ARG40 | CYT12 | vdW | 34.13 |
| ARG40 | URA11 | vdW | 31.34 |

| **AR** | **SLNCR20** | **Interaction** | **Occupancy (%)** |
|---|---|---|---|
| ARG9 | CYT8 | vdW | 26.35 |
| TYR11 | CYT18 | π-π Stacking | 35.73 |
| TYR11 | CYT18 | H-BOND | 28.14 |
| TYR11 | URA7 | vdW | 26.75 |
| TYR11 | URA7 | H-BOND | 25.35 |
| TYR11 | CYT18 | vdW | 24.35 |
| ARG13 | CYT17 | CATION-π | 23.35 |
| ARG13 | CYT17 | H-BOND | 22.55 |
| PRO14 | CYT18 | vdW | 26.95 |
| GLN35 | CYT17 | H-BOND | 27.74 |
| ASN36 | CYT17 | vdW | 35.53 |
| ASN36 | CYT17 | H-BOND | 32.14 |
| PRO39 | CYT17 | vdW | 33.93 |

| | | | |
|---|---|---|---|
| ARG40 | CYT15 | vdW | 57.09 |
| ARG40 | CYT17 | vdW | 52.69 |
| ARG40 | CYT15 | CATION-π | 45.51 |
| ARG40 | CYT17 | CATION-π | 38.52 |
| ARG40 | CYT17 | H-BOND | 36.13 |
| ARG40 | ADE16 | vdW | 23.95 |

**Supplementary Table S4.** Residue-level protein-RNA interactions were identified in the MD trajectories after applying a 20% occupancy cutoff. For each interaction, the protein residue, RNA nucleotide, interaction type, and occupancy percentage are reported for the $AR^{1-100}$-HOTAIR-20nt and $AR^{1-100}$-SLNCR1-20nt complexes.

| AR | HOTAIR20 | Interaction | Occupancy (%) |
|---|---|---|---|
| ARG9 | URA14 | vdW | 22.19 |
| ARG13 | CYT2 | vdW | 43.54 |
| ARG13 | GUA15 | H-BOND | 20.79 |
| ARG13 | GUA16 | H-BOND | 20.79 |
| ARG20 | ADE1 | vdW | 34.83 |
| GLN24 | ADE1 | H-BOND | 30.34 |
| GLN24 | ADE1 | vdW | 24.72 |
| GLN28 | ADE1 | vdW | 25.28 |
| ARG31 | URA11 | CATION-π | 75.84 |
| ARG31 | CYT12 | vdW | 34.27 |
| ARG31 | URA11 | vdW | 31.74 |
| ARG31 | CYT13 | vdW | 21.91 |
| ARG31 | CYT12 | H-BOND | 20.51 |
| ILE34 | CYT12 | vdW | 24.44 |
| GLN35 | CYT13 | vdW | 26.4 |
| ARG40 | CYT13 | CATION-π | 25.84 |

| AR | SLNCR20 | Interaction | Occupancy (%) |
|---|---|---|---|
| ARG13 | URA14 | vdW | 32.71 |

| | | | |
|---|---|---|---|
| ARG13 | URA14 | H-BOND | 31.91 |
| ARG13 | URA14 | CATION-π | 20.48 |
| PRO14 | GUA15 | H-BOND | 68.35 |
| PRO14 | GUA15 | vdW | 32.45 |
| PRO15 | GUA15 | H-BOND | 42.55 |
| PRO15 | GUA15 | vdW | 23.14 |
| SER16 | GUA15 | vdW | 23.4 |
| LYS17 | GUA15 | vdW | 28.46 |
| TYR19 | CYT5 | H-BOND | 59.84 |
| TYR19 | CYT5 | π-π Stacking | 49.47 |
| TYR19 | CYT5 | vdW | 25.8 |
| ARG20 | CYT7 | vdW | 29.52 |
| GLY21 | CYT6 | H-BOND | 68.35 |
| GLY21 | CYT6 | vdW | 42.29 |
| GLN24 | CYT6 | vdW | 30.05 |
| GLN24 | CYT6 | H-BOND | 21.01 |
| ARG31 | CYT13 | vdW | 25.27 |
| ARG31 | CYT12 | vdW | 21.01 |
| GLU32 | CYT13 | vdW | 24.2 |
| GLN35 | CYT12 | vdW | 21.54 |
| ARG40 | CYT12 | H-BOND | 62.77 |
| ARG40 | CYT13 | CATION-π | 54.52 |
| ARG40 | CYT12 | vdW | 41.22 |
| ARG40 | CYT13 | vdW | 29.79 |

**Supplementary Table S5.** Residue-level protein-RNA interactions identified in the HADDOCK-derived MD refinement trajectories after applying a 20% occupancy cutoff. For each interaction, the protein residue, RNA nucleotide, interaction type, and occupancy percentage are reported for the top-ranked HADDOCK models of the AR$^{1\text{-}100}$-HOTAIR-20nt and AR$^{1\text{-}100}$-SLNCR1-20nt complexes after MD refinement.

### Supplementary Figures

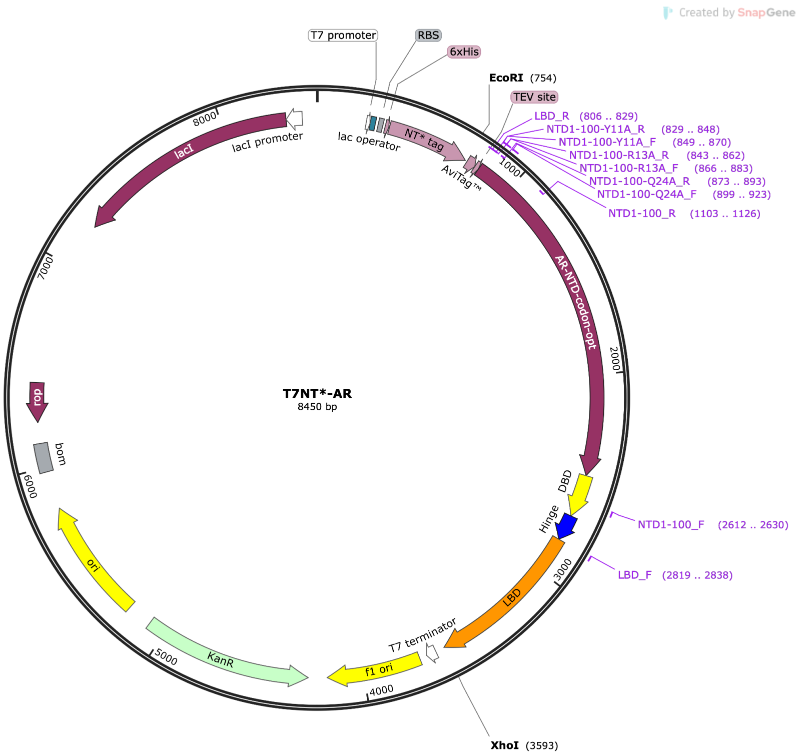


**Supplementary Figure S1. Vector map of the T7NT*$_{MaSp}$-AR plasmid used for bacterial expression of recombinant AR constructs.** A TEV cleavage site was inserted between the NT*MaSp solubility tag and the AR coding sequence, allowing the solubility tag to be removed during protein purification. The NT*MaSp tag has an N-terminal 6×His tag, allowing the cleaved tag to be separated from purified AR using reverse HisTrap purification. The vector map was exported from SnapGene software.

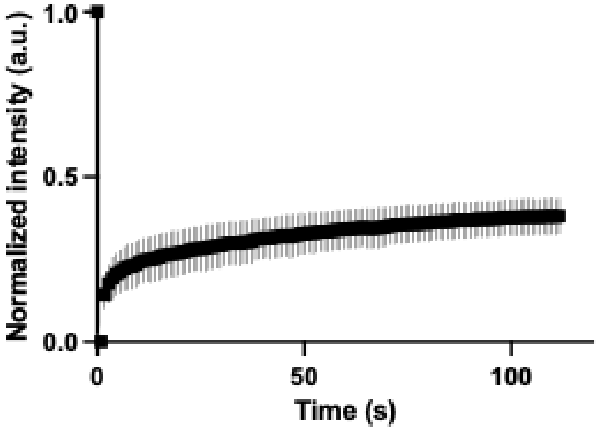


**Supplementary Figure S2. FRAP analysis of droplets formed by 20 μM AR$^{NTD}$ and 20 μM AR$^{LBD}$** (scale bar = 1 μm, 3 technical replicates).

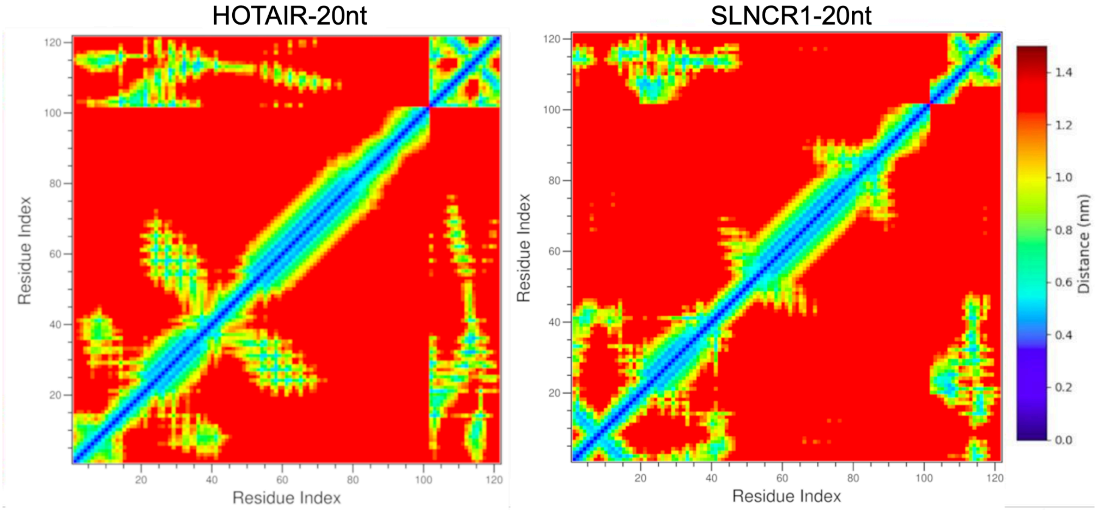


**Supplementary Figure S3. Mean minimum-distance contact maps obtained from the HADDOCK-derived MD refinement trajectories of AR$^{1\text{-}100}$ in complex with HOTAIR-20nt (left) and SLNCR1-20nt (right).** Each heatmap reports, for every residue pair, the mean of the minimum protein-RNA distances computed over the trajectory. Lower values (blue) indicate closer, more persistent contacts, whereas higher values (yellow to red) indicate greater separation. The HADDOCK-refined trajectories reproduce the main interaction regions identified in the unbiased simulations, including the compact interaction region centered on residues 13-31 for HOTAIR-20nt and the broader multi-site interaction pattern observed for SLNCR1-20nt.

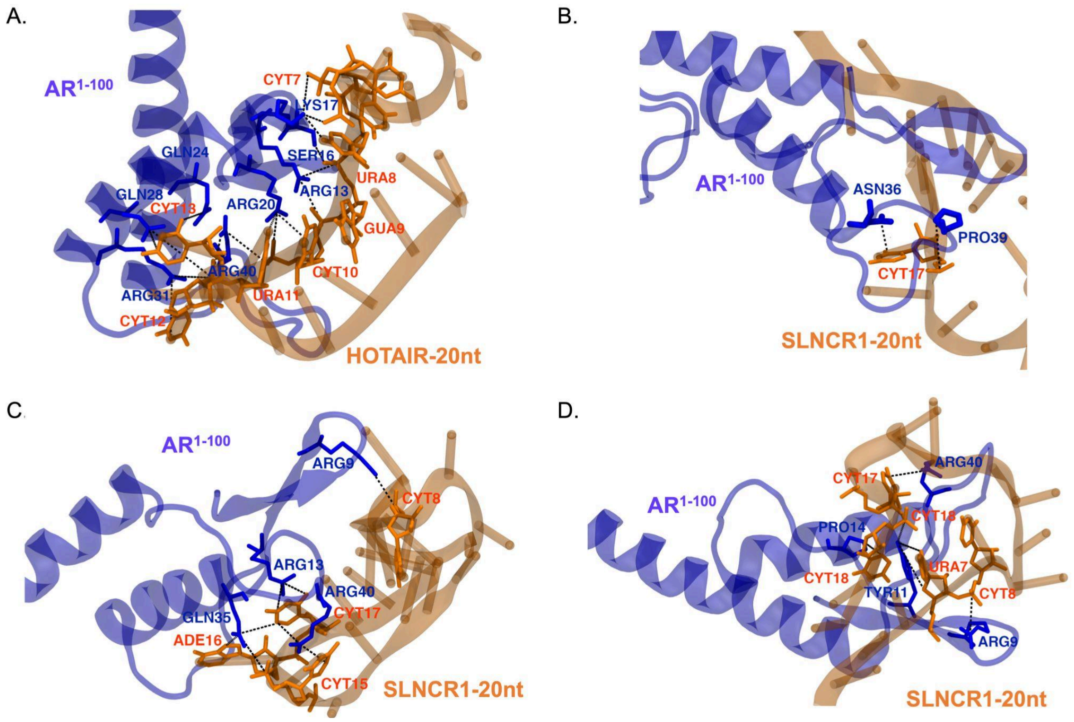

**Supplementary Figure S4. Representative MD snapshots of AR$^{1-100}$-RNA interactions.** Representative snapshots showing the highlighted residue-nucleotide contacts observed for A) HOTAIR-20nt at 2000 ns, showing contacts involving Arg31-Cyt12, Gln28-Cyt13, Gln24-Cyt13, Arg40-Cyt12/Ura11, Arg20-Cyt10/Ura11, Arg13-Gua9/Ura8, Ser16-Gua9, and Lys17-Cyt7/Ura8. B-D) three SLNCR1-20nt trajectories: SLNCR1-20nt-157 at 1657 ns, SLNCR1-20nt-349 at 1849 ns, and SLNCR1-20nt-498 at 1998 ns, with the indicated residue-nucleotide contacts highlighted in each snapshot.
